# An RNA dynamic ensemble at atomic resolution

**DOI:** 10.1101/2020.05.13.092981

**Authors:** Honglue Shi, Atul Rangadurai, Hala Abou Assi, Rohit Roy, David A. Case, Daniel Herschlag, Joseph D. Yesselman, Hashim M. Al-Hashimi

## Abstract

Biomolecules do not fold into a single 3D structure but rather form dynamic ensembles of many inter-converting conformations^1^. Knowledge of dynamic ensembles is key for understanding how biomolecules fold and function, and for rationally manipulating their activities in drug discovery and synthetic biology^2–4^. However, solving dynamic ensembles of biomolecules at atomic resolution is a major challenge in structural biology because the information required to specify the position of all atoms in thousands of conformations in an ensemble far exceeds the information content of experimental measurements. Here we addressed the data gap and dramatically simplified and accelerated RNA ensemble determination by using structure prediction tools that leverage the growing database of RNA structures to generate a conformational library. Library refinement with NMR residual dipolar couplings enabled determination of an atomic-resolution ensemble for HIV-1 TAR as confirmed by quantum-mechanical calculations of NMR chemical shifts, comparison to a crystal structure of a substate, and through the successful redistribution of the ensemble by design using atomic mutagenesis. The ensemble provides an unprecedented view of how bulge residues cooperatively flip out and undergo sugar repuckering to allow the adjoining helices to stack. The generality of this approach will make determination of atomic-resolution RNA ensembles routine.

## Main Text

The functions of many regulatory RNAs crucially depend on changes in three-dimensional (3D) structure that occur in response to a diverse array of cellular inputs, including the binding of small-molecule ligands^5, 6^, proteins^7^, epitranscriptomic modifications^8^, mutations^9^, and even to nascent RNA elongation during transcription^10, 11^. Initially described as changes from one 3D structure to another, these transitions are now better understood as changes in dynamic ensembles of many inter-converting conformations, from one conformational distribution to another^4, 12^.

To deeply understand RNAs at a level that ultimately makes it possible to rationally manipulate their behavior in drug discovery and synthetic biology, we need an ability to determine their dynamic ensembles at atomic resolution. This however presents a significant challenge to current biophysical techniques. The information required to specify the position of all atoms in thousands of conformations in an ensemble far exceeds the information content of experimental measurements. NMR spectroscopy is a rich source of ensemble-averaged measurements, and in combination with computational modelling, has been applied with success to determine ensembles of proteins at atomic resolution^13–15^. In contrast, fewer atomic-level experimental measurements are typically available for characterizing RNA ensembles. Moreover, while computational modeling methods such as molecular dynamics (MD) simulations are needed to address the experimental data gap^16^, nucleic acid force fields remain underdeveloped relative to proteins, and MD simulations of RNAs often poorly predict experimental data even for simple motifs^17, 18^. Consequently, relative to proteins, there is a greater danger of over-fitting an RNA ensemble, and assessing ensemble accuracy is more difficult.

To address the data gap, as well as dramatically simplify and accelerate RNA ensemble determination, we took advantage of structure prediction tools that leverage the growing database of RNA structures to directly generate a conformational library from a secondary structure that broadly samples energetically favorable 3D conformations. We used Fragment Assembly of RNA with Full-Atom Refinement (FARFAR)^19^, given its high performance in extensive tests of blind prediction of 3D RNA structure^20^. We then determined RNA ensembles by using NMR residual dipolar coupling (RDC) data^21–23^ to guide selection of conformers from the FARFAR library^18, 24, 25^.

We tested our approach on the transactivation response element (TAR) (Fig. 1a) from HIV-1^26, 27^, which has served as a model system for bulge motifs. Bulges are one of the most common RNA secondary structural elements that act as dynamic joints connecting helical elements, enabling their relative orientation to change adaptively during folding and function^18, 26–28^. We used FARFAR to directly generate a conformational library (*N* = 10,000) from an input TAR secondary structure only constraining Watson-Crick base pairs (bps) inferred by NMR, and assuming an idealized A-form geometry^29^ for these bps while predicting the structure for all remaining nucleotides (Methods). We then tested and optimized the FARFAR-library using a rich dataset of four independent RDCs (∼8 RDCs per nucleotide) previously measured for four TAR molecules variably elongated to modulate alignment relative to the NMR magnetic field^18, 25^.

**Fig. 1.**
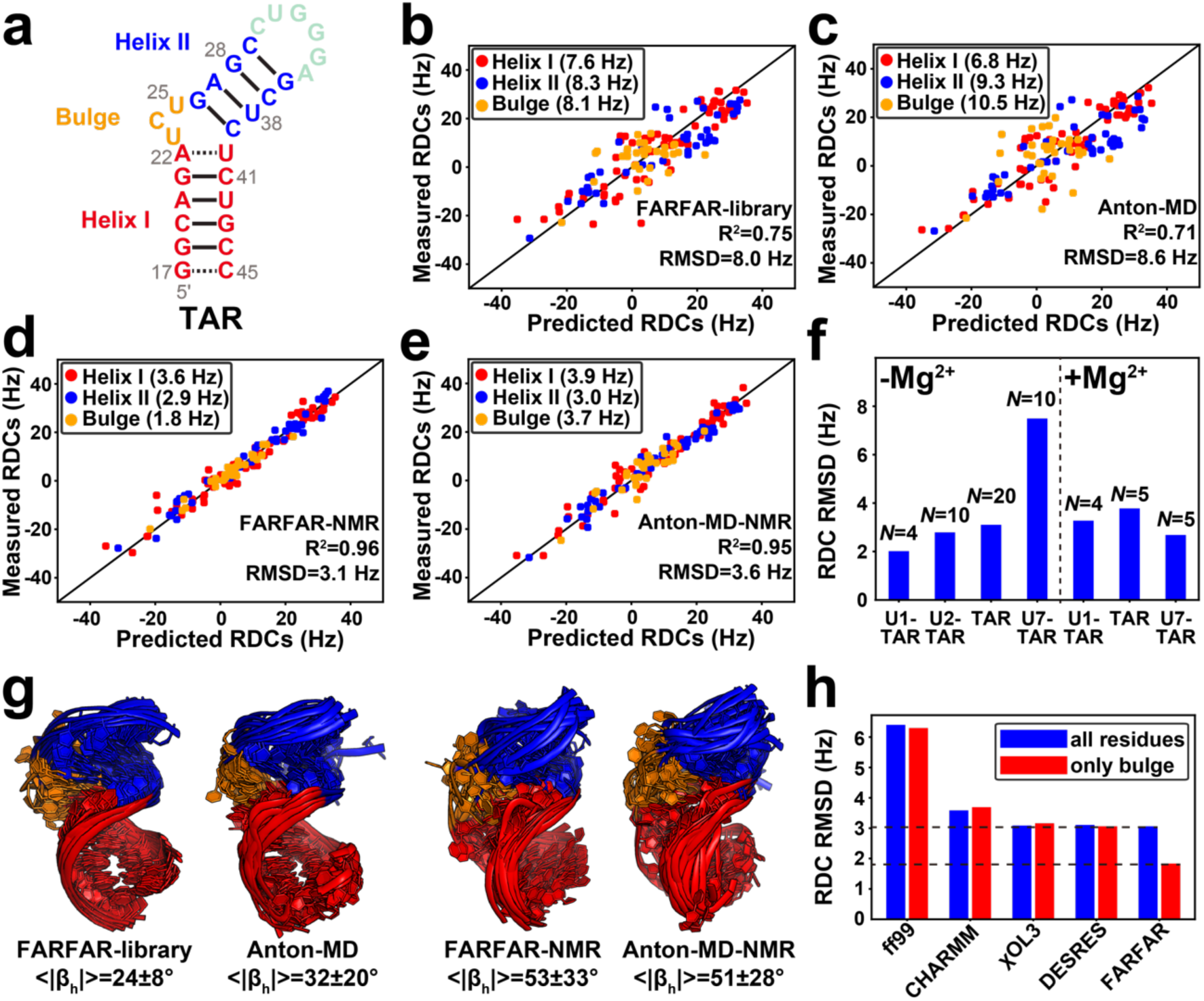
Using FARFAR-NMR to determine a TAR ensemble. (a) Secondary structure of TAR. (b-e) Comparison between measured and predicted TAR RDCs for the (b) FARFAR-library (*N* = 10,000), (c) Anton-MD library (*N* = 10,000), (d) FARFAR-NMR (*N* = 20) and (e) Anton-MD-NMR (*N* = 20). Values in parentheses denote the RDC RMSD. RDCs are color-coded according to the structural elements in Fig. 1a. (f) RDC RMSD for ensembles of TAR variants with ensemble size *N* obtained using FARFAR-NMR. (g) Structural overlay of the TAR ensembles (*N* = 20) (Methods). |β_h_| is the absolute magnitude of the bend angle. (h) Comparison of RDC RMSD over all (blue) or bulge (red) residues for ensembles generated using different MD force fields (ff99^30^, CHARMM36^31^, ff99bsc0χOL3^32^ and modified ff14 DESRES^33^) and FARFAR.

Intriguingly, the FARFAR-library showed better agreement (RMSD 8.0 versus 8.6 Hz) with the RDCs (Fig. 1b) as compared to a previously reported^18^ TAR library (Anton-MD) generated by subjecting an experimentally determined NOE-based NMR structure of TAR (PDB ID 1ANR)^26^ to MD simulations with the CHARMM36 force field^31^ (Fig. 1c). An optimized ensemble with *N* = 20 conformers (Extended Data Fig. 1b) was generated (Methods) by using the RDC agreement to guide selection of conformers from the FARFAR-library (Fig. 1d)^18, 24^. The optimized FARFAR-NMR ensemble also better predicted the RDCs (RMSD 3.1 Hz) relative to the optimized Anton-MD-NMR ensemble (RMSD 3.6 Hz) obtained using a similar procedure and the Anton-MD pool (Fig. 1e and Extended Data Fig. 1b). Cross validation^18, 34^ showed that the improved RDC agreement was not due to over-fitting (Extended Data Fig. 1c). Similar RDC agreement (Fig. 1f) was obtained when using FARFAR-NMR to generate ensembles for TAR in the presence of Mg^2+^, and three additional TAR mutants^28^ containing one (U1-TAR), two (U2-TAR), and seven (U7-TAR) bulge nucleotides in the presence and absence of Mg^2+^ (Extended Data Fig. 2-3 and Supplementary Discussion), establishing the generality of the approach.

The improved agreement observed with the FARFAR-NMR ensemble was surprising given that the global inter-helical distribution^35^ of the Anton-MD-NMR TAR ensemble has been independently validated via X-ray scattering interferometry (XSI) ^36^. Indeed, the FARFAR-NMR and Anton-MD-NMR ensembles show comparable agreement with the RDCs measured for helical bps (Fig. 1d, e) and the two ensembles sample similar inter-helical orientational distributions (Fig. 1g and Extended Data Fig. 4a-d).

Rather, the improved agreement was primarily driven by the RDCs measured in the locally more flexible bulge residues. For the FARFAR-NMR ensemble, the RDC RMSD (1.8 Hz) for bulge residues (Fig. 1d) was within experimental error (∼2 Hz), but was substantially higher (3.7 Hz) for the Anton-MD-NMR ensemble (Fig. 1e). The bulge RDC RMSD could not be improved by running MD simulations using different force fields (Fig. 1h), indicating that the improved conformational sampling in the FARFAR-library makes it possible to surpass the accuracy with which the TAR bulge could be described using conventional MD simulations. This improved performance is particularly noteworthy when considering that determining a high-resolution structure and running MD can take several months and often years whereas the FARFAR-library is generated within 24 hours running on 100 cores in parallel.

To further evaluate the accuracy of the FARFAR-NMR TAR ensemble, we substantially expanded the breadth and depth of atomic-level experimental data that can be brought to bear when evaluating the accuracy of RNA ensembles by predicting ensemble-averaged ^1^H, ^13^C, and ^15^N chemical shifts (∼15 chemical shifts per nucleotide) using quantum-mechanical AF-QM/MM calculations^37, 38^. Although rich in structural information that is complimentary to that obtained from RDCs, the agreement between measured chemical shifts and values predicted from crystal structures of nucleic acids has traditionally been poor^37, 39^. We recently showed in studies of DNA duplexes that at least some of this disagreement originates from neglecting ensemble averaging when predicting ^13^C chemical shifts^38^. This revelation and the improved FARFAR-NMR TAR ensemble led us to test this approach on flexible RNAs, and to extend its reach by incorporating ^1^H and ^15^N shifts in addition to ^13^C chemical shifts (Fig. 2a).

**Fig. 2.**
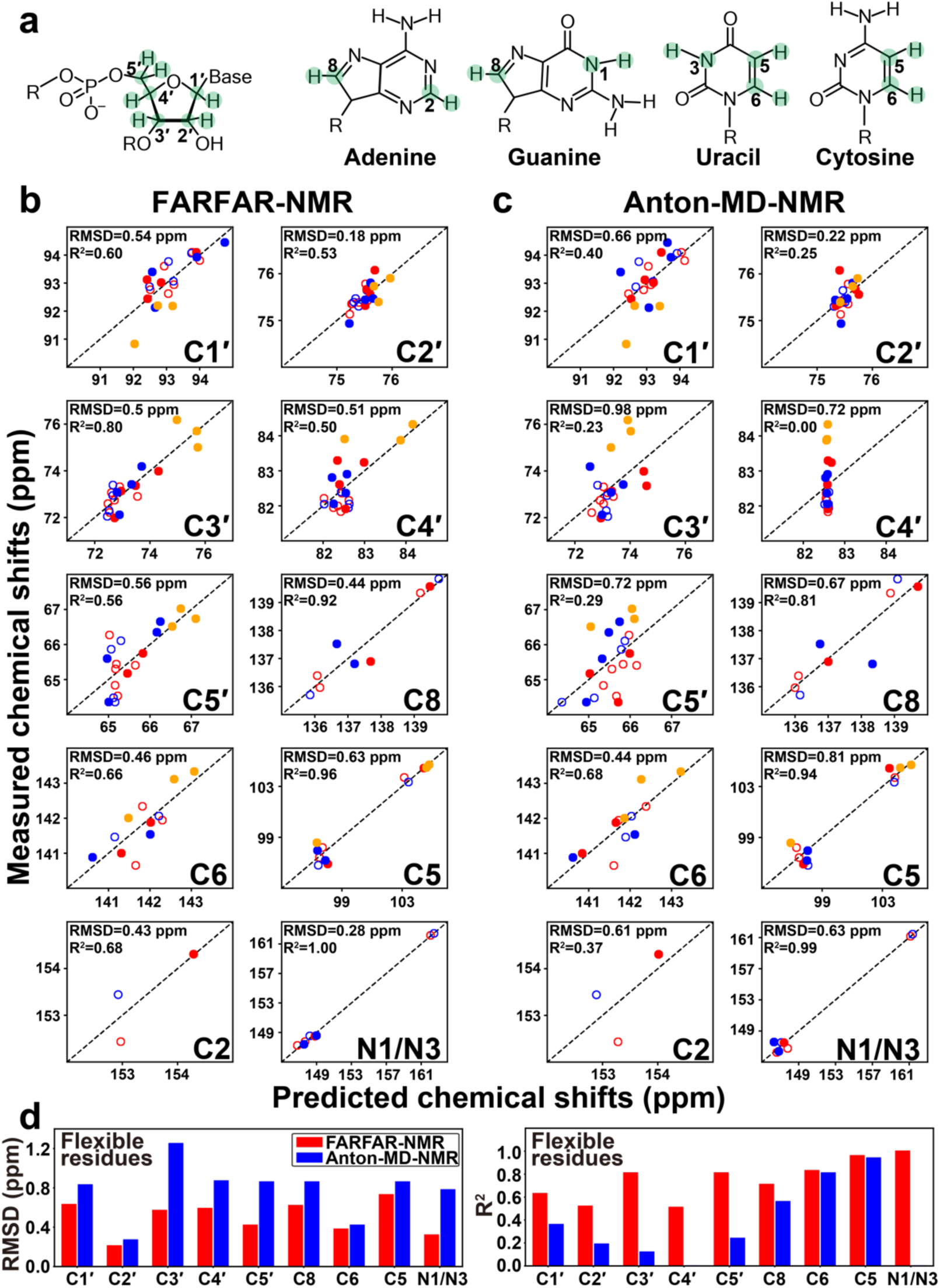
Evaluating TAR ensembles using chemical shifts. (a) Chemical structures of the sugar and base moieties with chemical shift probes used to test ensemble accuracy highlighted in green. (b-c) Comparison of measured and predicted ^13^C/^15^N chemical shifts for (b) FARFAR-NMR (*N* = 20) (c) Anton-MD-NMR (*N* = 20) by AF-QM/MM (Methods). Values are color-coded according to the structural elements in Fig. 1a. Chemical shifts for central Watson-Crick bps within A-form helices (C19-G43, A20-U42, G21-C41, A27-U38, G28-C37) are denoted using open circles. A correction was applied to the predicted chemical shifts (Methods) as described previously^38^. For ^1^H chemical shifts see Extended Data Fig. 5b, d. (d) Comparison of RMSD (left) and R^2^ (right) between measured and predicted ^13^C/^15^N chemical shifts for flexible residues (U23, C24, U25, A22-U40, G26-C39, C29-G36, G18-C44) for FARFAR-NMR (red) and Anton-MD-NMR (blue).

Remarkably, good agreement was observed between the measured ^1^H, ^13^C, and ^15^N base and sugar chemical shifts and values back-calculated for the FARFAR-library (Extended Data Fig. 5 and Supplementary Discussion). The agreement improved substantially for the FARFAR-NMR ensemble following RDC optimization for 48% of the atom-types analyzed, while the agreement was unaffected for the remaining atom-types (Fig. 2b and Extended Data Fig. 5). The agreement deteriorated substantially for any one conformer member of the ensemble, underscoring the critical importance of ensemble-averaging (Extended Data Fig. 6). The agreement with an independent set of measurements not used in ensemble determination that is highly sensitive to various structural features of the base, sugar, and backbone strongly suggests that FARFAR-NMR describes the TAR ensemble with atomic accuracy (< 2 Å, Methods). In sharp contrast, the agreement was much weaker for the Anton-MD-NMR ensemble relative to FARFAR-NMR for 90% of the atom-types, particularly for sugar chemical shifts, some of which show no apparent correlation even following RDC optimization (Fig. 2c, d and Extended Data Fig. 5). These results establish the utility of chemical shifts in RNA ensemble determination and also show that the improved accuracy of the FARFAR-NMR ensemble relative to Anton-MD-NMR is even greater than alluded to by the RDC data.

We compared the FARFAR-NMR and Anton-MD-NMR ensembles (Fig. 3a) to better understand the features responsible for the more accurate ensemble description of the TAR bulge. Approximately 75% of the conformers in the FARFAR-NMR ensemble have the canonical UCU bulge, while the remaining 25% have a non-canonical AUC bulge that forms through a single nucleotide register shift (Extended Data Fig. 7a). In contrast, conformers in the Anton-MD-NMR ensemble sample a broader set of junction topologies some of which diverge from the NMR-derived Watson-Crick pairing (Extended Data Fig. 7b and Supplementary Discussion), and this could contribute to the poor agreement observed with the imino ^15^N/^1^H chemical shifts (Fig. 2c and Extended Data Fig. 5).

**Fig. 3.**
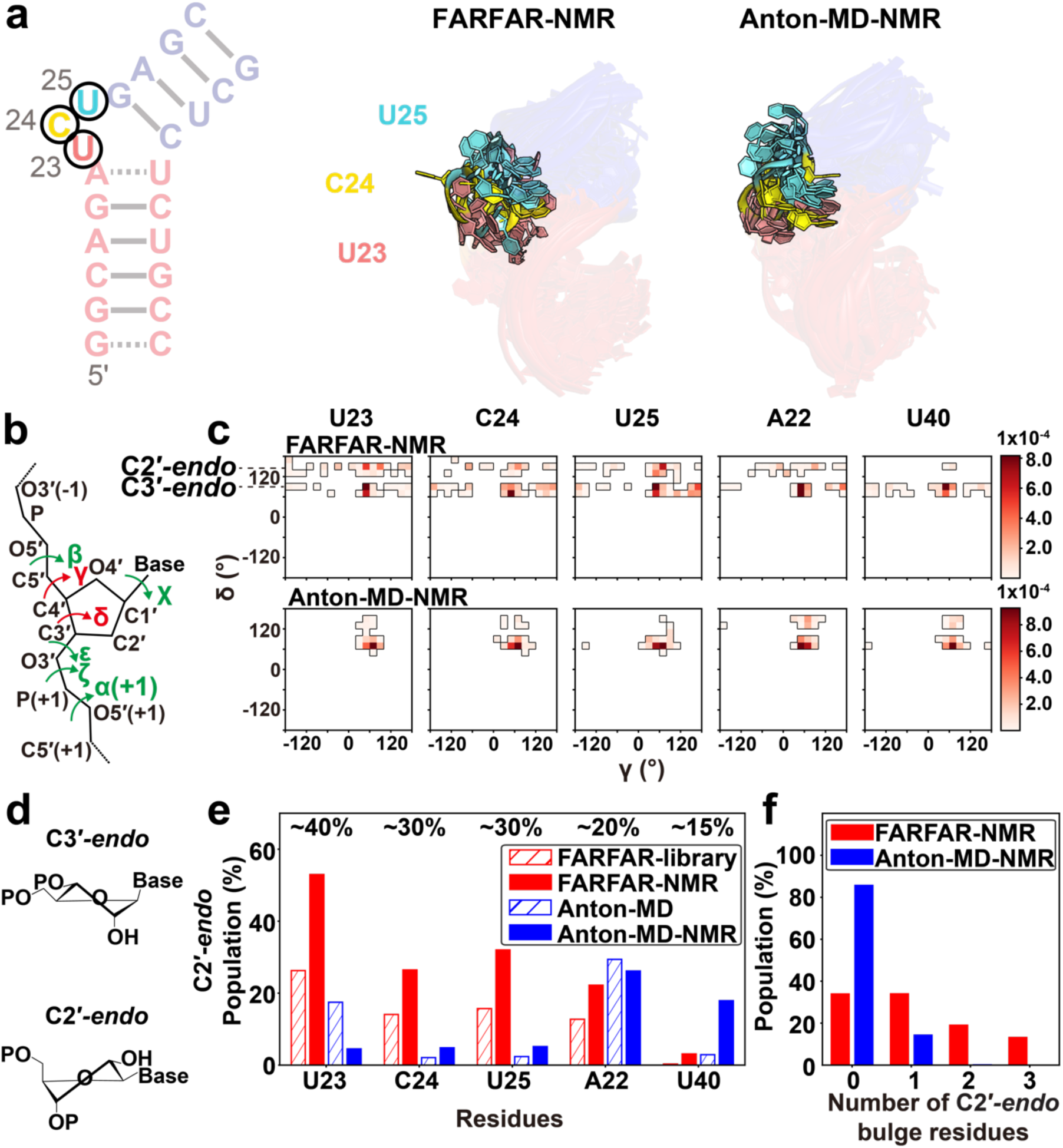
FARFAR-NMR ensemble more broadly samples non-canonical sugar-backbone torsion angles relative to Anton-MD-NMR. (a) Overlay of the dynamic ensembles of the TAR bulge. (b) RNA backbone torsion angles exhibiting different and similar distributions between Anton-MD-NMR and FARFAR-NMR are colored red and green, respectively (Methods). (c) 2D density maps of δ versus γ comparing Anton-MD-NMR and FARFAR-NMR ensembles (*N* = 2,000) for bulge residues as well as A22 and U40. The bin width is 20°. (d) Structure of the ribose moiety in C3′-*endo* and C2′-*endo* conformations. (e) Population of C2′-*endo* pucker at bulge residues as well as A22 and U40 in the FARFAR-library (*N* = 10,000, red open), FARFAR-NMR (*N* = 2,000, red fill), Anton-MD library (*N* = 10,000, blue open) and Anton-MD-NMR (*N* = 2,000, blue fill). Experimental estimates of the C2′-*endo* population based on ^13^C chemical shifts are indicated above the bars (Methods). (f) The population of conformers in the ensemble as a function of the number of C2′-*endo* bulge residues for FARFAR-NMR (red, *N* = 2,000) and Anton-MD-NMR (blue, *N* = 2,000).

In addition, despite having a narrower range of junction topologies, the sugar pucker distribution (defined by angle δ) for bulge residues in the FARFAR-NMR sample a broader range (Fig. 3b, c), are substantially enriched in the non-canonical C2′-*endo* conformation (Fig. 3d) relative to the Anton-MD-NMR ensemble, and are in better agreement with the NMR-derived^40^ sugar puckers (Fig. 3e and Extended Data Fig. 8a). Moreover, ∼15% of the conformers in the FARFAR-NMR ensemble had all three bulge residues simultaneously in the C2′-*endo* conformation but none did in the Anton-MD-NMR ensemble (Fig. 3f). Excluding all conformers containing C2′-*endo* sugar puckers in U23, C24, U25, A22 or U40 from the FARFAR-library diminished the RDC agreement to a level similar to the Anton-MD derived ensembles again confirming that the improved agreement is not due to over-fitting of the data (Extended Data Fig. 8b, c). A similar behavior was observed for the backbone torsion angle γ (Fig. 3b), which shows greater sampling of non-*gauche*+ angles in the FARFAR relative to the Anton-derived ensembles (Fig. 3c and Extended Data Fig. 8b, d). This can account for the better agreement observed for sugar chemical shifts^40^ in the FARFAR-NMR versus Anton-MD-NMR ensembles (Fig. 2b-d).

What causes conformers in the FARFAR-NMR ensemble to more greatly sample non-canonical sugar-backbone conformations at the bulge relative to conformers in the Anton-MD-NMR ensemble? Unpaired pyrimidine RNA nucleotides unconstrained by other interactions are enriched in the C2′-*endo* conformation (C2′-*endo*:C3′-*endo* is 40:60)^41^ and C3′-*endo* becomes the predominant sugar-pucker when the nucleotides form bps or stack intra-helically^40^. Indeed, ∼80% of the residues with C2′-*endo* sugar puckers were extra-helical in the FARFAR-NMR ensemble (Fig. 4a and Extended Data Fig. 7a). Moreover, for linear inter-helical conformations, there was a strong preference to have all three bulge residues simultaneously flip out (Fig. 4b) and adopt conformations enriched in the C2′-*endo* sugar pucker (Fig. 4c); the bulge residues flip out to allow the two helices to coaxially stack (Supplementary Video 1 and Extended Data Fig. 7a). Such coaxial conformations have previously been hypothesized to exist within the TAR ensemble based on the Mg^2+^ dependence of the inter-helical ensemble^28^ and a crystal structure of Ca^2+^ bound TAR^42^.

**Fig. 4.**
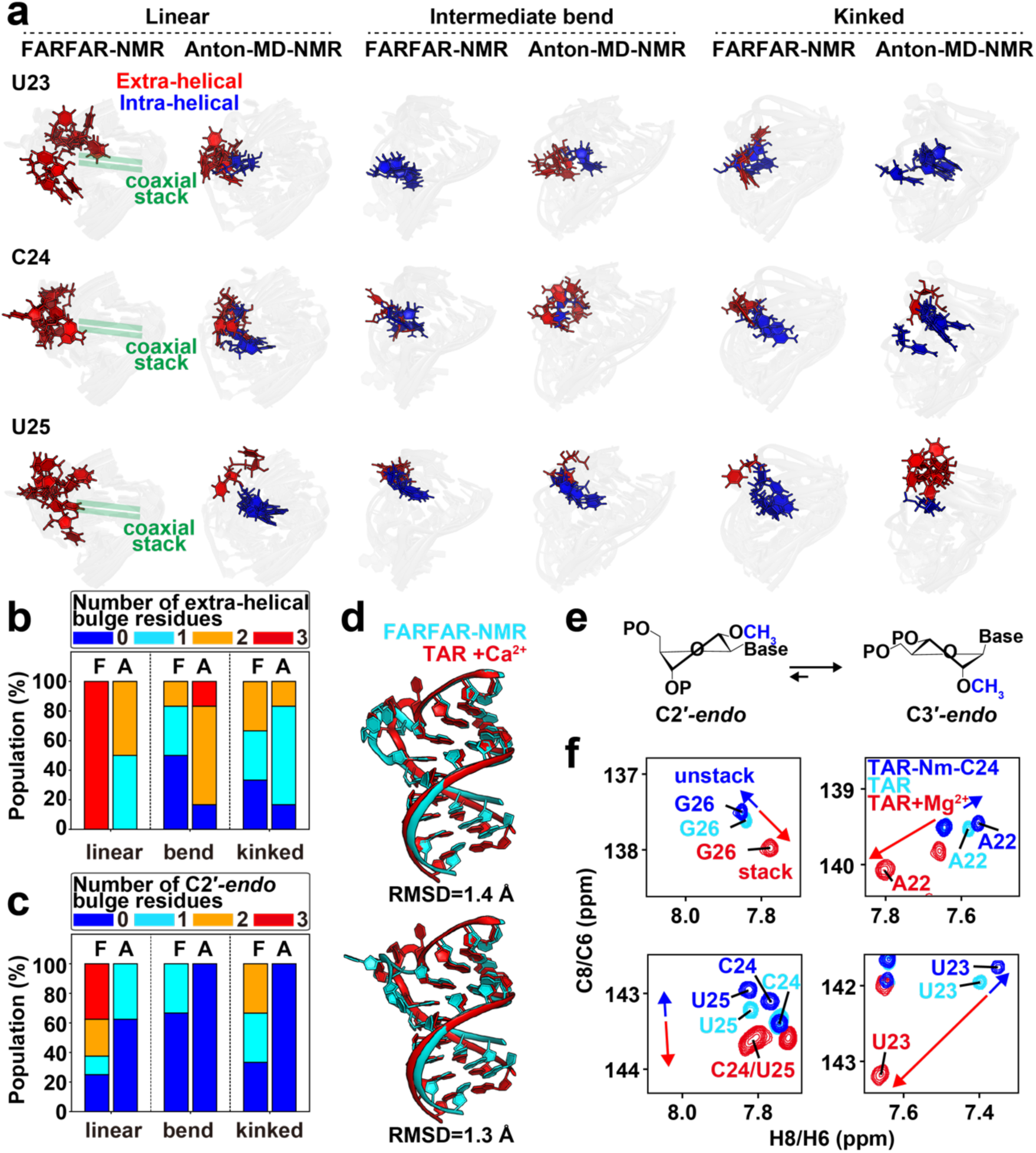
Cooperative extra-helical flipping and sugar repuckering of bulge residues are coupled to coaxial stacking. (a) Overlay of conformers showing motions in bulge residues for linear (|β_h_| < 45°), intermediate bend (45° < |β_h_| < 70°) and kinked (|β_h_| > 70°) inter-helical conformations in the FARFAR-NMR and Anton-MD-NMR ensembles. (b-c) The fractional populations of conformers with (b) extra-helical and (c) C2′-*endo* bulge residues (color-coded) as a function of bending angle in the FARFAR-NMR and Anton-MD-NMR ensembles. F and A denote FARFAR-NMR and Anton-MD-NMR, respectively. (d) Comparison of two coaxially stacked FARFAR-NMR conformers with the crystal structure of Ca^2+^ bound TAR (PDB: 397D)^42^. (e) Nm shifts sugar pucker equilibrium towards C3′-*endo*. (f) Overlay of 2D [^13^C, ^1^H] NMR HSQC spectra of the aromatic spins for TAR-Nm-C24 without Mg^2+^ (blue, inducing unstacking), TAR without Mg^2+^ (cyan) and TAR with Mg^2+^ (red, inducing stacking).

Strikingly, two of the coaxial conformations in the FARFAR-NMR ensemble superimpose with the TAR crystal structure with a heavy-atom RMSD of 1.3-1.4 Å (Fig. 4d). Thus, the TAR crystal structure captures substates of the ensemble in solution. In sharp contrast, none of the conformations in the Anton-MD-NMR ensemble had the two helices coaxially stacked, and not a single conformer had all three bulge residues flipped out and having the C2′-*endo* sugar pucker (Fig. 4a-c and Extended Data Fig. 7b). The FARFAR-NMR ensemble (Fig. 4a and Supplementary Video 1) further shows that all three bulge residues simultaneously flip out with high cooperativity (∼2.3 kcal/mol, see methods) most likely because this then permits favorable coaxial stacking of the helices (Fig. 4a, Extended Data Fig. 7a and Supplementary Video 1). This can explain prior results^43^ showing that a TAR mutation that promotes inter-helical stacking results in a predominantly coaxially stacked conformation in which all three bulge residues are flipped out.

Finally, we put key atomic features of the FARFAR-NMR ensemble to a test by rationally redistributing the conformer populations using atomic mutagenesis. The ensemble shows that the kinked unstacked conformations are enriched in bulge residues that have the C3′-*endo* sugar pucker relative to the coaxial stacked conformations (Fig. 4c). To test this feature of the ensemble, we incorporated 2′-O-Methyl (Nm) modifications (Fig. 4e) at U23 or C24 to bias the sugar pucker at these positions toward C3′-*endo* by ∼0.2 kcal/mol^44^. Indeed, methylating either U23 or C24 resulted in chemical shift perturbations in 2D NMR spectra of TAR (Fig. 4f and Extended Data Fig. 9) throughout the bulge and neighboring residues that are specifically directed towards the chemical shifts of the unstacked conformation^28^, as expected for a cooperative redistribution in favor of the kinked conformation.

In conclusion, FARFAR-NMR lays the foundation for a new paradigm for RNA ensemble determination by combining reliably measurable NMR data with 3D structure prediction. The approach can immediately be applied to render many existing NMR structures of RNAs that were determined using conventional approaches into dynamic ensembles, and their accuracies can be tested using the available chemical shift data. The approach is general, rapid, and can also incorporate other sources of experimental data. Given its ease of implementation and higher throughput, FARFAR-NMR has the potential to unleash the ensemble description of RNAs to all corners of biology, from which a deeper and broader understanding of folding and function will undoubtedly emerge.

## Methods

### Generating ensembles using FARFAR

FARFAR^19^ is implemented as the *rna_denovo* program in Rosetta Software Suite. FARFAR requires as input the RNA sequence, which can be constrained by an optional secondary structure. The input secondary structures for TAR were derived based on the imino ^1^H resonances in NMR spectra. Therefore, Watson-Crick base pairing was imposed for G18-C44, C19-G43, A20-U42, G21-C41, G26-C39, A27-U38, G28-C37 and C29-G36 whereas nucleotides in the apical loop and bulge, and the terminal G17-C45 bp were set to be unconstrained. Given the indirect NMR observation of the A22-U40 bp in U2-TAR by H6(C5)NN experiment as reported previously^28^, two sets of FARFAR simulations were conducted for each TAR variant with A22-U40 either paired or unpaired. Interestingly, with the exception of U2-TAR, constraining A22-U40 when generating FARFAR-libraries did not improve the agreement with the RDCs for the TAR and other variants. This highlights the importance of differentiating between well-formed versus labile bps, an important advantage offered by NMR as compared to other methods for determining secondary structure such as chemical probing. Thus, all the FARFAR-libraries in this study correspond to those in which the A22-U40 base pair is not constrained, except for the U2-TAR ensemble in the absence of Mg^2+^. To significantly reduce the run time, we modeled the helices of TAR and its bulge variants as static idealized A-form helices shown previously to satisfy NMR RDC data in RNA^28, 29^. *rna_helix.py* is a python wrapper for the Rosetta executable *rna_helix*. *rna_helix.py* is available in $ROSETTA/tools/rna_tools/bin, where $ROSETTA is the Rosetta installation path.

~~~
rna_helix.py –seq gcag cugc –resnum 18-21 41-44 –o helix1.pdb
rna_helix.py –seq gagc gcuc –resnum 26-29 36-39 –o helix2.pdb
~~~

in which RNA sequences of both strands as well as their residue indices are required as input. While generating FARFAR-library assuming idealized A-form helices, we then executed *rna_denovo* with the following command:

~~~
rna_denovo –nstruct 100 –s helix_1.pdb helix_2.pdb –fasta input.fasta –secstruct_file input.secstruct –minimize_rna true
~~~

where *`–nstruct`* is the number of models per run which is 100 in the current study, *`–fasta`* is the path of a fasta file containing the RNA sequence, *`–secstruct`* is the path of a file (input.secstruct) containing the RNA secondary structure in dot-bracket notation, *`–minimize_rna true`* minimizes the RNA after fragment assembly, and *`–s`* specifies the path to the pdb files that contain static structures of our helices we do not wish to generate via fragment assembly to save computation (the fasta, secstruct files as well as example commands can be found in Supplementary Table 2-5).

The entire procedure was repeated 100-200 times and 10,000 structures were randomly selected from the entire resulting output with Rosetta energy units < 0 to remove models that potentially may have chain breaks and severe steric clash which do not satisfy the RDCs (Extended Data Fig. 1a), to generate the FARFAR-library (*N* = 10,000).

### Molecular dynamics (MD) simulations

Simulations of HIV-1 TAR starting from the PDB structure 1ANR using the CHARMM36 force field^31^ were performed as described previously^18^. MD simulations of TAR starting from the PDB structure 1ANR using the ff99 force field^30^ with and without χ_OL3_ corrections for RNA^32^ were performed using periodic boundary conditions as implemented in the AMBER MD simulation package^45^. All starting structures were solvated using a truncated octahedral box of TIP3P^46^ water molecules with box size chosen such that the boundary was at least 11 Å away from any of the RNA atoms for all simulations with the ff99 force field with χ_OL3_ corrections. For the TAR simulations with the ff99 force field without χ_OL3_ corrections, the system was solvated in a truncated octahedral box of SPC/E water molecules such that the boundary was at least 15.4 Å away from any of the RNA atoms. Na^+^ ions treated using the Joung-Cheatham parameters^47^ were then added to neutralize the charge of the system in all cases. The system was then energy minimized in two stages with the solute being fixed (with a restraint of 500 kcal/mol/Å^2^) during the first stage. Equilibration and production runs (1 μs) were then performed as described previously^48^. Simulations of TAR starting from PDB 1ANR using the DESRES force field^33^ were performed using the GROMACS MD simulation package^49, 50^. DESRES force field files for GROMACS were obtained from a port by Giovanni Bussi (https://github.com/srnas/ff/tree/desres). The starting structure (PDB 1ANR) was solvated using a rhombic dodecahedral box of TIP4P-D^51^ water molecules, with box size chosen such that the boundary was at least 10 Å away from any of the RNA atoms. Na^+^ ions treated using the parameters from MacKerell *et al*.^52^ were then added to neutralize the charge of the system. After energy minimization without restraints, the system was then gradually heated to a temperature of to 298 K using a modified Berendsen thermostat with a stochastic term^53, 54^ (τ = 0.1 ps), under constant volume conditions for 100 ps with harmonic restraints on the solute (1000 kJ/mol/nm^2^). The system was then allowed to equilibrate for 100 ps under constant pressure (1 bar), using the Parinello-Rahman barostat^55, 56^ (τ = 2 ps) and temperature (at 298 K, using a modified Berensen thermostat, τ = 0.1 ps) conditions, with harmonic restraints on the solute (1000 kJ/mol/nm^2^). This was followed by NPT equilibration for 30 ns without harmonic restraints, following by a production run of 1 μs. A non-bonded cutoff of 9 Å was used for treating short range non-bonded interactions while the Particle Mesh Ewald method^57^ was used to treat long range electrostatic interactions. Covalent bonds involving hydrogen were constrained using the LINCS algorithm^58^ to enable the use of a 2 fs timestep. A set of evenly (5 ps) spaced snapshots was used for subsequent analysis of all simulations using the CPPTRAJ suite of programs^59^.

Additional MD simulations using the ff99 force field with χ_OL3_ corrections were also performed to assess the extent to which sampling of sugar puckers in the bulge could be influenced by changes in the starting structures used for the simulations (Extended Data Fig. 10). Two starting structures were derived from PDB 1ANR with the sugar puckers of U23 (TAR^U23C2′-*endo*^) and U25 (TAR^U25C2′-*endo*^) individually switched from C3′-*endo* to C2′-*endo*, and another was a conformer from the FARFAR-NMR ensemble (TAR^FARFAR^) in which the sugar puckers of U23, C24 and U25 were all C2′-*endo*. The starting structures for the TAR^U23C2′-*endo*^ and TAR^U25C2′-*endo*^ simulations were generated by superposition of a C2′-*endo* uridine nucleotide onto U23/U25 in 1ANR using the uridine base atoms, and replacing the sugar-backbone atoms with those of the superimposed C2′-*endo* uridine. For TAR^U23C2′-*endo*^ and TAR^U25C2′-*endo*^, only the backbone of the C2′-*endo* uridine was fixed during energy minimization, while for TAR^FARFAR^, all the heavy atoms were fixed during minimization.

### NMR residual dipolar coupling (RDC) data

The RDC data (sugar C1′-H1′/C2′-H2′/C3′-H3′/C4′-H4′ and base C8-H8/C6-H6/C2-H2/C5-H5/N1-H1/N3-H3) used in the TAR ensemble determination were reported previously^18, 25, 60^ and are summarized in Supplementary Table 1. The raw data can also be downloaded from https://sites.duke.edu/alhashimilab/resources/. RDCs were measured using the same buffer conditions (15 mM sodium phosphate, 25 mM NaCl, 0.1 mM EDTA, with/without 3 mM Mg^2+^ at pH ∼ 6.4) at 25°C with similar RNA concentrations (∼1.0 mM) and slightly different (6-22 mg/mL) Pf1 phage concentrations ^18^.

### RDC calculations

Ensemble-averaged RDCs were calculated by computing the RDCs for each conformer in an ensemble using the program PALES^61^. PALES computes RDCs based on global molecular shape. The RDC were computed using a cylindrical wall model using the following command:

~~~
pales –pdb input.pdb –inD input_rdc.tab –outD output_rdc.tab –H –pf1 –wv 0.022
~~~

where *`–pdb`* is the path of an input PDB file (input.pdb), *`–inD`* is the path of an input data file indicating the bond vectors for which RDC should be computed (input_rdc.tab), *`–outD`* is the path of the output file containing all the calculated RDCs, *`–H`* means selecting all atoms including proton and *`–pf1 –wv`* specifies the pf1 effective concentration (0.022g/mL) assuming a rod liquid crystal model.

The RDC values were the averaged over all conformers in an ensemble assuming that they are equiprobable. Individual scaling factors were applied on the predicted RDCs of each construct to account for the difference of alignment magnitude in part arising due to differences in phage concentrations in the experiments, as described previously^18^. The elongated constructs were elongated *in silico* using an idealized A-form geometry prior to RDC calculation as described previously^18^.

The TAR apical loop was modeled using the wild-type CUGGGA loop^18^. As reported previously for the Anton-MD-NMR ensemble^18^, replacing the loop with the UUCG loop used to measure RDCs^62^ minimally impacted the RDC agreement for the FARFAR ensembles (Extended Data Fig. 1e, f). This is consistent with prior NMR studies showing that the apical loop replacement minimally impacts the dynamics of the bulge^18, 63^.

### Sample and Select (SAS)

We used the SAS approach^24^ to generate ensembles from a structural pool that best satisfied the measured RDCs. Briefly, a simulated annealing Monte Carlo sampling scheme was used to select an ensemble that minimizes the cost function depicting the differences between the measured and predicted RDCs:

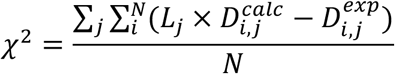

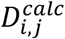 and 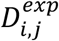 are the calculated and measured RDCs, respectively, of the *ith* bond vector measured on the *jth* TAR construct, *L_j_* is the overall scaling factor of alignment magnitude for construct *j*, and *N* is the total number of bond vectors. The initial effective temperature for simulation annealing was 100 and decreased by a factor of 0.9 in every step for a total of 5×10^5^ steps. A series of SAS runs were performed varying the ensemble size from *N* = 1 to an ensemble size in which the RDC RMSD reaches a plateau (Extended Data Fig. 1b, 2). The resultant ensemble size for different bulge variants were: *N* = 4 (U1-TAR, no Mg^2+^), *N* = 10 (U2-TAR, no Mg^2+^), *N* = 20 (TAR, no Mg^2+^), *N* = 10 (U7-TAR, no Mg^2+^), *N* = 4 (U1-TAR, with Mg^2+^), *N* = 5 (TAR, with Mg^2+^) and *N* = 5 (U7-TAR, with Mg^2+^). To analyze distributions of structural parameters (Fig. 3 and Extended Data Fig. 3, 4), we also generated larger sized ensembles by running SAS multiple times to ensure the total size *N* = 2,000 for all systems.

The SAS analysis excluded RDCs from the flexible terminal G17-C45 bp as well as those of the C29-G36 bp flanking the apical loop, given differences between the apical loop sequences used to measure RDCs (wild-type or UUCG) and to model (wild-type) TAR. Note that RDCs from G17-C45 and C29-G36 bp were included in the prior study^18^ and this explains the small differences in RDC RMSD for the two Anton-MD derived ensembles relative to that reported earlier.

### Cross-validation analysis

Cross-validation was performed using two approaches as described previously^18^. In one approach (inactive random, Extended Data Fig. 1c, d), 10% of the RDC data was randomly removed and SAS was used to generate an ensemble. The RMSD between measured and predicted RDCs was then computed for the left out RDC data. This procedure was repeated 10 times and the final RDC RMSD was averaged over all 10 independent runs. For TAR (absence of Mg^2+^) where we have RDCs measured on four constructs, a second mode (inactive media, Extended Data Fig. 1c, d) of cross-validation was also performed, in which the RDC dataset of each construct was left out individually before running SAS. The RMSD between measured and predicted RDCs was then computed for the left out RDC dataset. The final RDC RMSD is averaged over the four iterations corresponding to leaving out each RDC dataset.

### NMR chemical shift data

The ^1^H, ^13^C and ^15^N chemical shift assignments of TAR have been published previously^40, 63^ and were compared to quantum mechanical chemical shift predictions. The numerical populations for C2′-*endo* shown in Fig. 3e were obtained based on the C1′ chemical shift assuming a linear dependence between the range 86 ppm (100 % C2′-*endo*) ^48^ and 94 ppm (100 % C3′-*endo*)^40^.

### Automated fragmentation quantum mechanics/ molecular mechanics (AF-QM/MM) chemical shift calculation

Chemical shift calculations were performed using a previously described fragmentation procedure^37^. Each RNA structure was subjected to 5 steps of conjugate gradient minimization with harmonic restraints of 2 kcal/mol-Å^2^ on all heavy atoms; this regularizes bond lengths and angles to minimize noise in the results that can arise from very small changes in these geometric parameters. Next, each structure was broken into “quantum” fragments centered on each nucleotide, containing 2-6 neighboring nucleotides, using a heavy-atom distance cutoff of 3.4 Å. The effects of RNA atoms outside the quantum region, and of water and ions in the solvent, were represented as point charges uniformly distributed on the molecular surface of the quantum region and resolved by fitting to Poisson−Boltzmann calculations using the “solinprot” program from the MEAD package^64, 65^. The quantum region was assigned a local dielectric ε of 1 (vacuum); the remaining RNA region had an ε of 4, and the solvent region an ε of 80. GIAO chemical shift calculations were carried out for each fragment, using version 5.0 of the demon-2k program^66^ using the OLYP functional^67^ with the pcSseg-1 (triple-ς plus polarization) basis set optimized for chemical shifts^68^ for the central nucleotide (whose results are reported here), and a DZVP basis for the remaining atoms. Reference shieldings were computed for tetramethylsilane (TMS) using the same functional and basis set. The ensembles of Anton-MD-NMR, FARFAR-NMR as well as a randomly selected Anton-MD and FARFAR-library of size *N* = 20 were examined. Note that R^2^ can be artificially low for spins with small ranges of chemical shifts (e.g. ∼1 ppm for C3′/C4′/C6 for central Watson-Crick bps).

### Ensemble analysis

The local backbone and sugar torsion angles (Fig. 3b) were calculated using X3DNA-DSSR^69^. The inter-helical Euler angles (α_h_, β_h_, γ_h_) were computed as described previously^35^ (Fig. 1g, Fig. 4b, c, Extended Data Fig. 3, 4, 7). Briefly, the upper helix from G26 to G28 and lower helix from C19 to G21 are aligned to idealized A-form helix, respectively. The sign of α_h_ and γ_h_ is inverted relative to previously reported values^18^ such that a positive and negative inter-helical twist angle (α_h_+γ_h_) corresponds to over- and under-twisting, respectively^70^.

Junctional topology (Extended Data Fig. 7) was defined as the base pairing mode which is detected by X3DNA-DSSR^69^. If two bases are forming a base pair with Leontis-Westhof (LW) classification as “cWW” (e.g. Watson-Crick base pair, Wobble base pair), a solid line is indicated between the two bases (Extended Data Fig. 7), whereas other LW classifications are indicated as a dash line (Extended Data Fig. 7).

TAR conformers were considered to be co-axially stacked when the bases comprising A22-U40 and G26-C39 or U25-U40, were stacked with each other, as defined by X3DNA-DSSR^69^.

The lower bound estimate of pairwise RMSD between two ensembles was defined as the following: Consider two ensembles A and B with size *N*_A_ and *N*_B_. For every conformer in ensemble A, we found the corresponding conformer in ensemble B that has the lowest pairwise RMSD. This procedure was repeated for every conformer in A to obtain a total of *N*_A_ RMSD values. We then took the root mean square of all these *N*_A_ RMSD values. The procedure was also repeated considering ensemble B. Then the minimum of the two root mean square values was selected as a lower bound of the similarity between the two ensembles. By this definition, Anton-MD-NMR and FARFAR-NMR ensembles differ by 3.7 Å while ensembles obtained from multiple independent FARFAR-NMR runs typically differ by 1.4 Å on average.

The cooperativity of flipping bulge nucleotides out was computed from the FARFAR ensemble as follows – For U23, C24 and U25, the probability of independently flipping out was computed as:

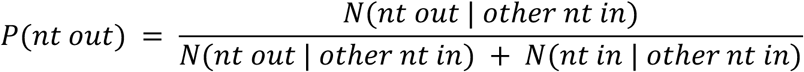

where *N(nt out | other nt in)* is the number of FARFAR-NMR conformers with the nucleotide (nt) of interest being flipped out with the other nucleotides (among U23, C24 and U25) being flipped in and *N(nt in | other nt in)* is the number of FARFAR-NMR conformers with U23, C24 and U25 flipped in. Flipping in and flipping out of U23, C24 and U25 were gauged by visual examination of the conformers (Extended Data Fig. 7). The probability of simultaneously flipping out U23, C24 and U25 without cooperativity as then computed as the product of the probabilities of independently flipping out each nucleotide as defined above i.e.,

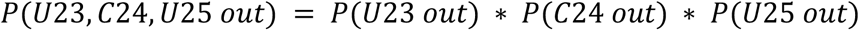

This was then compared to the observed probability of U23, C24 and U25 being flipped out *P(U23, C24, U25 out obs)* which was computed as the fraction of FARFAR-NMR conformers with U23, C24 and U25 flipped out (8/20). Cooperativity was defined as:

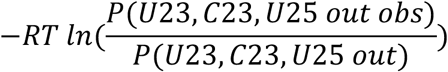

where *R* is the universal gas constant, *T* is temperature in Kelvin (298 K).

### NMR sample preparation

TAR, TAR-Nm-U23, and TAR-Nm-C24 RNA samples were synthesized using a MerMade 6 Oligo Synthesizer (BioAutomation) via solid-phase synthesis using standard phosphoramidite chemistry and deprotection protocols^71^. 2′-TBDMS protected phosphoramidites (ChemGenes) and 1 µmol standard synthesis columns (1000 Å) were used. The final 5′-DMT (4,4′-dimethoxytrityl) was removed during the synthesis for DMT-off deprotection and PAGE purification. Removal of nucleobase and phosphate protecting groups, and cleavage from the 1 µmol columns was achieved using 1 ml of 30% ammonium hydroxide and 30% methylamine (1:1) followed by 2-hour incubation at room temperature. The solution was then air-dried and dissolved in 100 µL DMSO and 125 µL TEA-3HF, followed by 2.5 h incubation at 65 °C following Glen Research protocols (https://www.glenresearch.com/reports/gr19-22) for 2′-O deprotection. The sample was then ethanol precipitated overnight, air dried, then dissolved in water for gel purification using a 20% (w/v) polyacrylamide gel with 8 M urea and 1 x Tris/borate/EDTA. The RNA was removed from the excised gel by electro-elution in 1 x Tris/acetic acid/EDTA followed by ethanol precipitation. The RNA was annealed in water at a concentration of 50 µM by heating at 95 °C for 5 min followed by cooling on ice for 60 min. It was then buffer exchanged using an Amicon Ultra-15 centrifugal filter (EMD Milipore) with a 3 kDa cutoff into NMR buffer (15mM sodium phosphate, 25 mM NaCl, 0.1mM EDTA) at pH 6.4. The final concentrations were: ∼0.8 mM for TAR, 2.5 mM TAR-Nm-U23 and 1.4 mM TAR-Nm-C24. TAR was also buffer exchanged into NMR buffer containing 3 mM Mg^2+^.

### NMR spectroscopy

All the NMR 2D HSQC experiments in this study were carried out on Bruker Avance III 600-MHz NMR spectrometer equipped with a triple-resonance cryogenic probed at 25°C. NMR Data were processed using NMRpipe^72^ and analyzed using SPARKY (T.D. Goddard and D.G. Kneller, SPARKY 3, University of California, San Francisco), respectively. The resonance assignments for Nm-modified TAR were obtained based on the previous reported assignments of TAR^63^ and further confirmed using 2D NOESY experiments.

## Acknowledgments

We thank members of the Al-Hashimi laboratory and Dr. Dawn Merriman (University of Dayton) for assistance and critical comments on the manuscript. We would like to thank Dr. Andrew Watkins (Stanford University) for advice about FARFAR.

## Funding

This work was supported by US National Institute for General Medical Sciences (1R01GM132899 to H.M.A. and D.H.), and US National Institute of Health (U54 AI150470 to H.M.A. and D.A.C.) and by Tobacco Settlement Fund (21-5734-0010 to J.D.Y.).

## Author contributions

H.S., D.H., J.D.Y. and H.M.A. conceived the project and experimental design. H.S. and J.D.Y. performed FARFAR calculations. A.R. performed MD simulations. H.S. performed sample and select and other ensemble analysis, with assistance from R.R. H.A.A. prepared NMR samples, performed NMR experiments and analyzed NMR data. D.A.C. performed QM/MM chemical shift calculation. H.M.A. and H.S. wrote the manuscript with critical input from A.R., D.H., D.A.C. and J.D.Y.

## Competing interests

H.M.A. is advisor to and holds an ownership interest in Nymirum, an RNA-based drug discovery company. The remaining authors declare no competing interests.

## Materials & Correspondence

Rosetta FARFAR commands as well as its required input files are included in Supplementary Materials. The raw RDC data can be downloaded from https://sites.duke.edu/alhashimilab/resources/. The AFNMR programs are available at https://github.com/dacase/afnmr. All other in-house Python scripts for SAS, structural analysis as well as other data analysis are available on request to the corresponding authors. The authors declare that the data supporting the findings are available within the article and its Supplementary Materials, or are available upon request to the corresponding authors.

## Extended Data Figures

**Extended Data Fig. 1.**
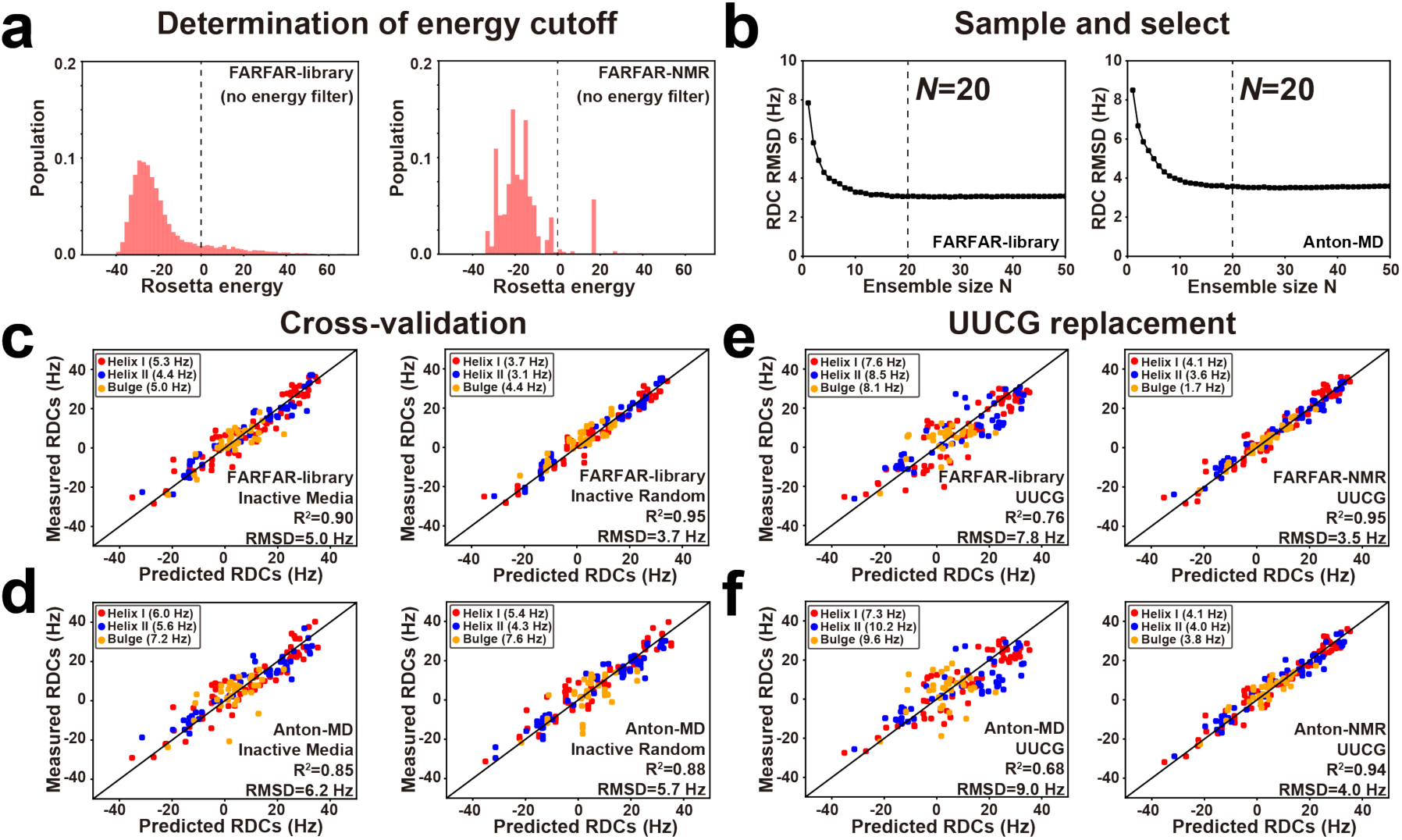
Optimizing TAR ensembles using SAS. (a) Determination of Rosetta energy cutoff. (left) Shown are the distribution of energies of a FARFAR-library (no energy filter) (*N* = 10,000) generated without pre-filtering based on Rosetta energy (i.e., not excluding structures with Rosetta energy > 0). (right) Distribution of no pre-filtering FARFAR-library in (left) after RDC selection (*N* = 2,000). The Rosetta energy = 0 is indicated using a dashed line. As structures with energy > 0 were predominantly excluded following SAS, only structures with energy < 0 were retained while generating the FARFAR-library in the main manuscript. (b) RDC RMSD as a function of ensemble size (*N*) during SAS for (left) the FARFAR-library and (right) Anton-MD. The chosen ensemble size *N* = 20 is indicated using a vertical dashed line. (c-d) Cross-validation analysis of TAR ensembles. Shown are comparison of measured and predicted RDCs obtained from cross-validation analysis on TAR ensembles using two modes: Inactive Media (left) and Inactive Random (right)(Methods) for (c) the FARFAR-library and (d) Anton-MD. (e-f) Generation of TAR ensembles using UUCG apical loop models. Comparison between measured and predicted RDCs for starting pools with *N* = 10,000 (left) and comparison between measured and predicted RDCs after SAS with *N* = 20 (right) from (e) FARFAR-library replacing wild-type CUGGGA loop with a UUCG loop^62^ and (f) Anton-MD replacing wild-type CUGGGA loop with a UUCG loop. Replacing the loop with the UUCG loop used to measure RDCs minimally impacted the RDC agreement for both FARFAR and Anton ensembles.

**Extended Data Fig. 2.**
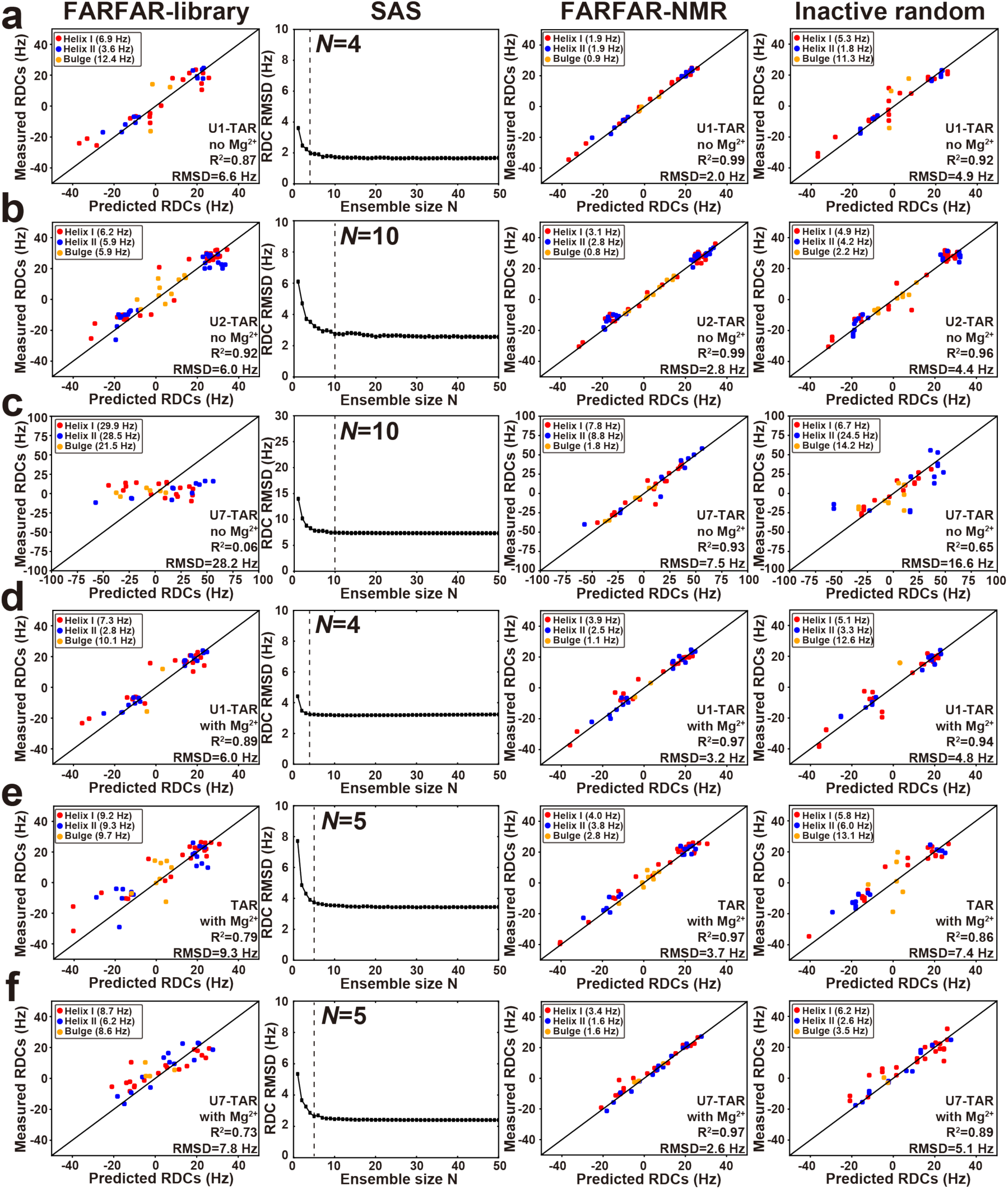
Optimizing ensembles of TAR and its variants. From left to right, comparison of measured and predicted RDCs for the FARFAR-library (*N* = 10,000), RDC RMSD as a function of ensemble size during SAS (ensemble size *N* chosen for the final ensembles is indicated as a vertical dashed line), comparison between the measured and predicted RDCs for the FARFAR-NMR ensembles after SAS, and inactive random cross-validation for (a) U1-TAR no Mg^2+^ (b) U2-TAR no Mg^2+^ (c) U7-TAR no Mg^2+^ (d) U1-TAR with Mg^2+^ (e) TAR with Mg^2+^ and (f) U7-TAR with Mg^2+^. RDC values are color coded according to structural element as defined in Fig. 1a and Extended Data Figure 3a.

**Extended Data Fig. 3.**
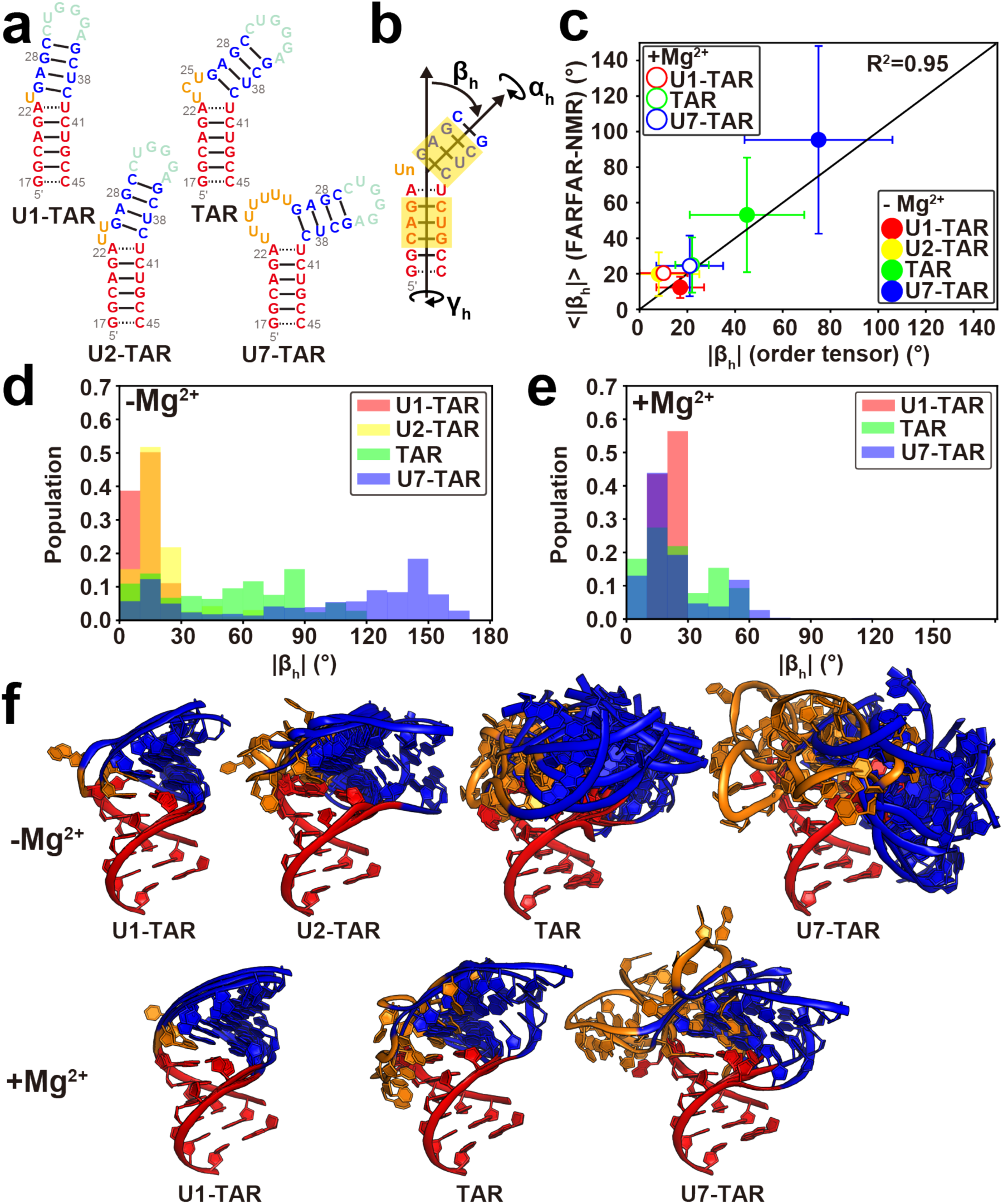
Dynamic ensembles of TAR and its bulge variants in the absence and presence of 3 mM Mg^2+^. (a) Secondary structure of TAR and its bulge variants (U1-TAR, U2-TAR and U7-TAR). (b) Definition of the three inter-helical Euler angles (α_h_, β_h_, γ_h_) describing the relative orientation of the two helices (Methods). The bps of the upper and lower helices used for defining the Euler angles are highlighted in yellow. (c) Comparison between the average bend angle (<|β_h_|>) and its standard deviation for the FARFAR-NMR (*N* = 2,000) derived ensembles with values obtained from an order tensor analysis of the RDCs^28^. The standard deviation in the order tensor analysis corresponds to half the cone radius angle assuming an isotropic model^73^. (d-e) Distributions of the inter-helical bend angle magnitude |β_h_| for each ensemble (*N* = 2,000) in the absence (d) and presence (e) of Mg^2+^. (f) Ensembles of TAR and its bulge variants in the absence (upper) and presence (lower) of 3 mM Mg^2+^. Motifs are color-coded as in (a).

**Extended Data Fig. 4.**
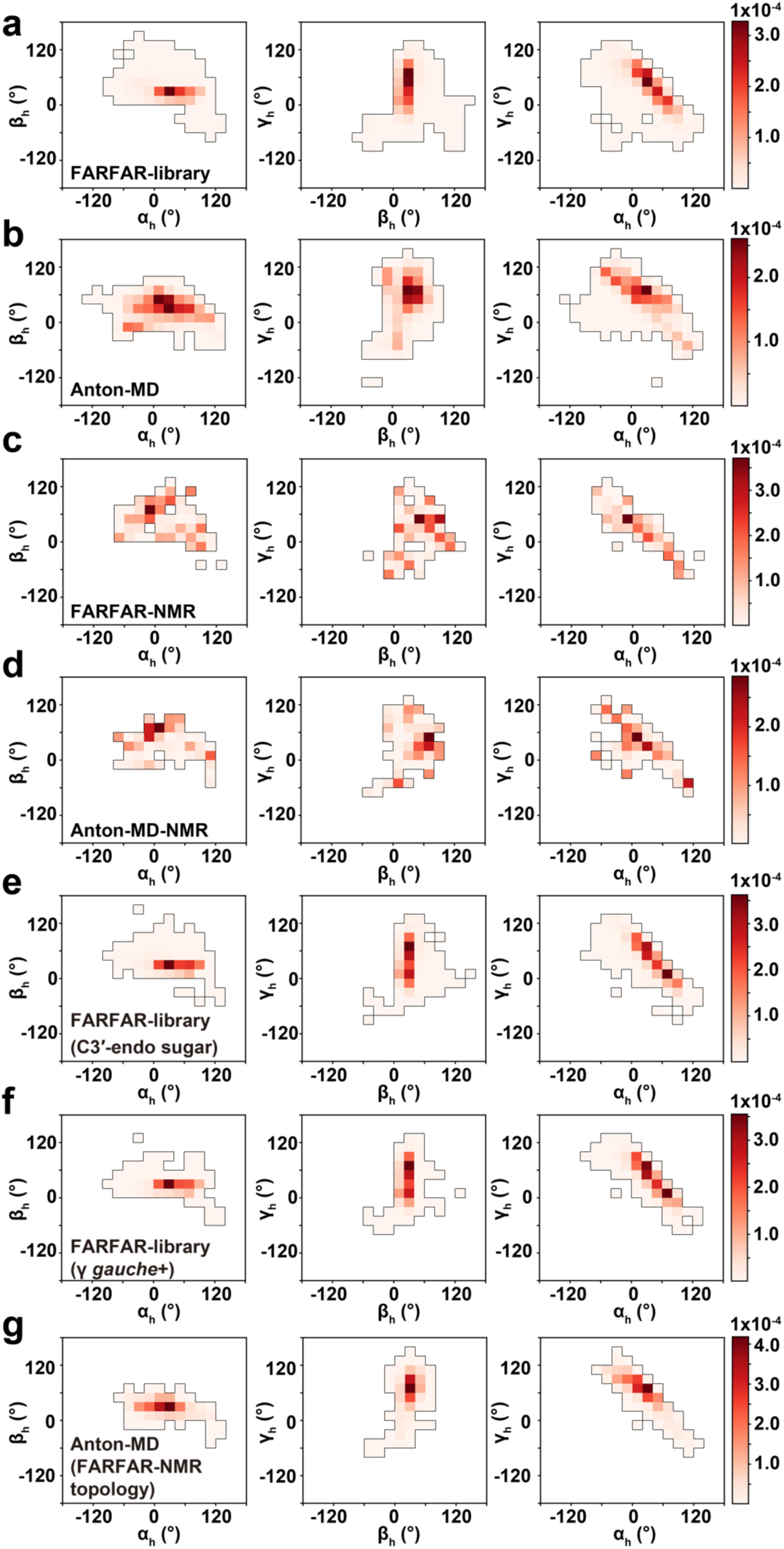
Inter-helical orientational distributions of FARFAR and Anton libraries and ensembles. The 2D density map of inter-helical Euler angle (α_h_, β_h_, γ_h_) for (a) FARFAR-library (*N* = 10,000), (b) Anton-MD (*N* = 10,000), (c) FARFAR-NMR (*N* = 2,000), (d) Anton-MD-NMR (*N* = 2,000), (e-f) two subsets of FARFAR-library: (e) one with U23, C24, U25, A22 and U40 constrained to be C3′-*endo* (*N* = 4,422), and (f) the other one with U23, C24, U25, A22 and U40 constrained to be only *gauche*+ γ(20°<γ<100°) (*N* = 3,145), (g) a subset of Anton-MD with conformers retaining the same junction topology as that in FARFAR-NMR (Methods) (*N* = 2,074). The color-scale for density is given on the right. In all cases, the bin width is 20°.

**Extended Data Fig. 5.**
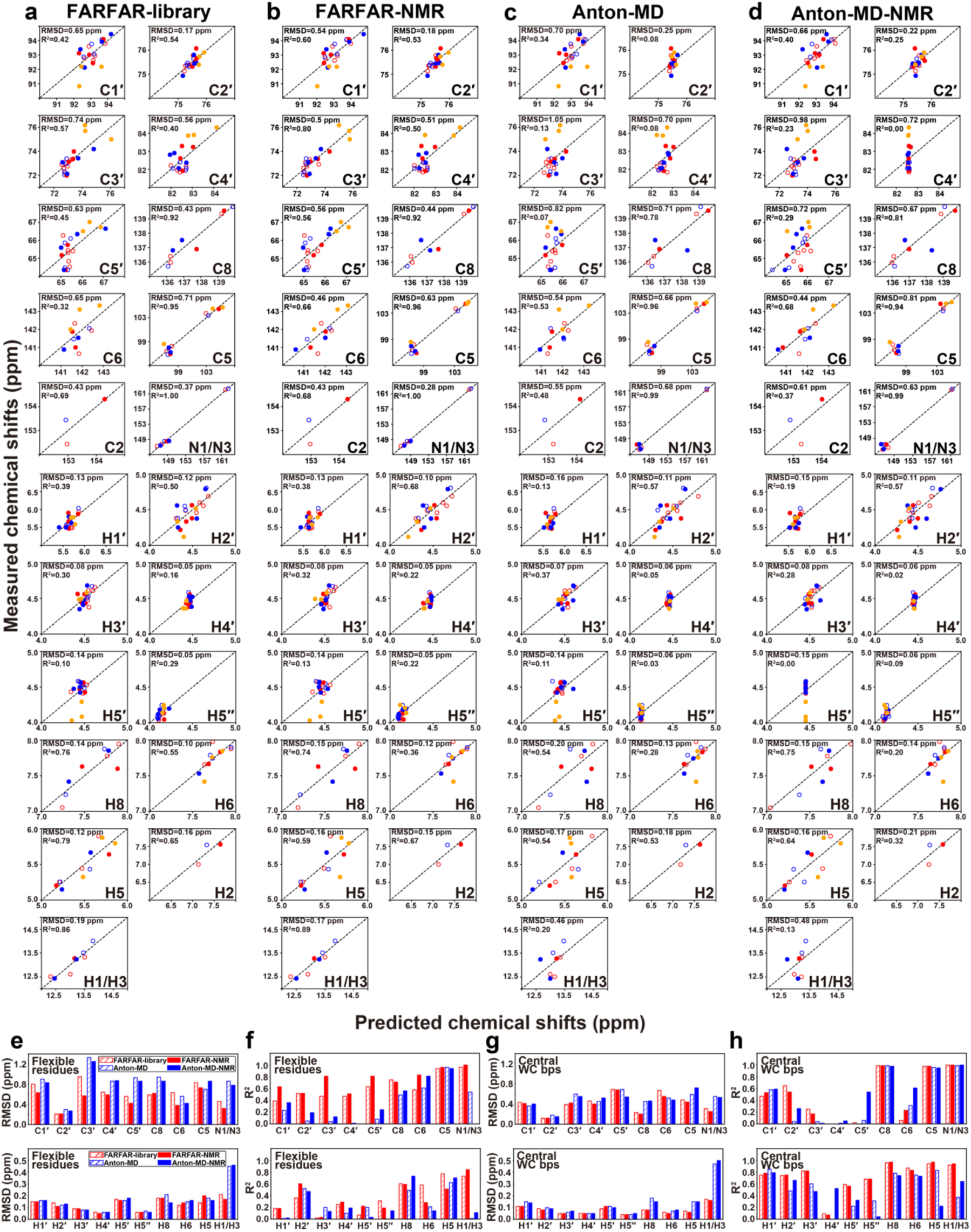
Evaluation of TAR libraries and ensembles via ^13^C, ^15^N and ^1^H chemical shifts. (a-d) Comparison of measured and predicted ^13^C, ^15^N and ^1^H chemical shifts for (a) FARFAR-library, (b) the FARFAR-NMR ensemble, (c) Anton-MD, and (d) the Anton-MD-NMR ensemble (*N* = 20 in all cases) by AF-QM/MM (Methods). Chemical shifts are color-coded according to the different structural elements (Fig. 1a). Chemical shifts for the central Watson-Crick bps within A-form helices (C19-G43, A20-U42, G21-C41, A27-U38, G28-C37) are denoted using open circles. A correction was applied to the predicted chemical shifts (Methods) as described previously^38^. (e-h) Comparison of RMSD and R^2^ between measured and predicted ^13^C/^15^N (top) and ^1^H (below) chemical shifts for (e-f) flexible residues (U23, C24, U25, A22-U40, G26-C39, C29-G36, G18-C44) and (g-h) central Watson-Crick bps for the FARFAR-library (red, open), FARFAR-NMR ensemble (*N* = 20) (red, fill), Anton-MD (blue, open) and the Anton-MD-NMR ensemble (*N* = 20) (blue, fill).

**Extended Data Fig. 6.**
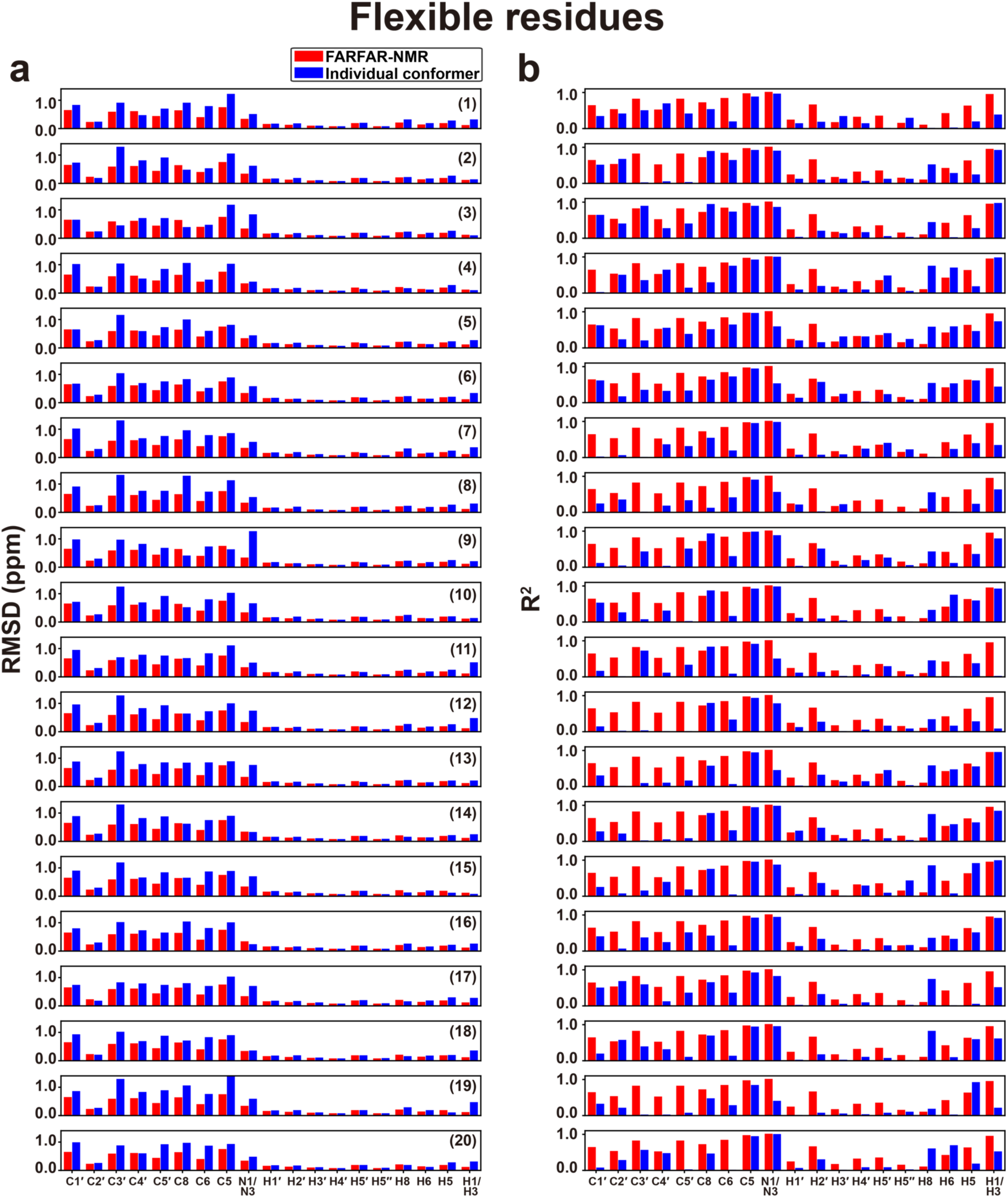
Comparison of agreement of measured and predicted chemical shifts between FARFAR-NMR ensemble and individual conformers. Bar plot of (a) RMSD and (b) R^2^ between measured and predicted chemical shifts for FARFAR-NMR ensemble (red) and its individual conformer (blue) for only flexible residues (U23, C24, U25, A22-U40, G26-C39, C29-G36, G18-C44). All the conformers in FARFAR-NMR ensemble (*N* = 20) are sorted in increasing order of the bend angle magnitude |β_h_|.

**Extended Data Fig. 7.**
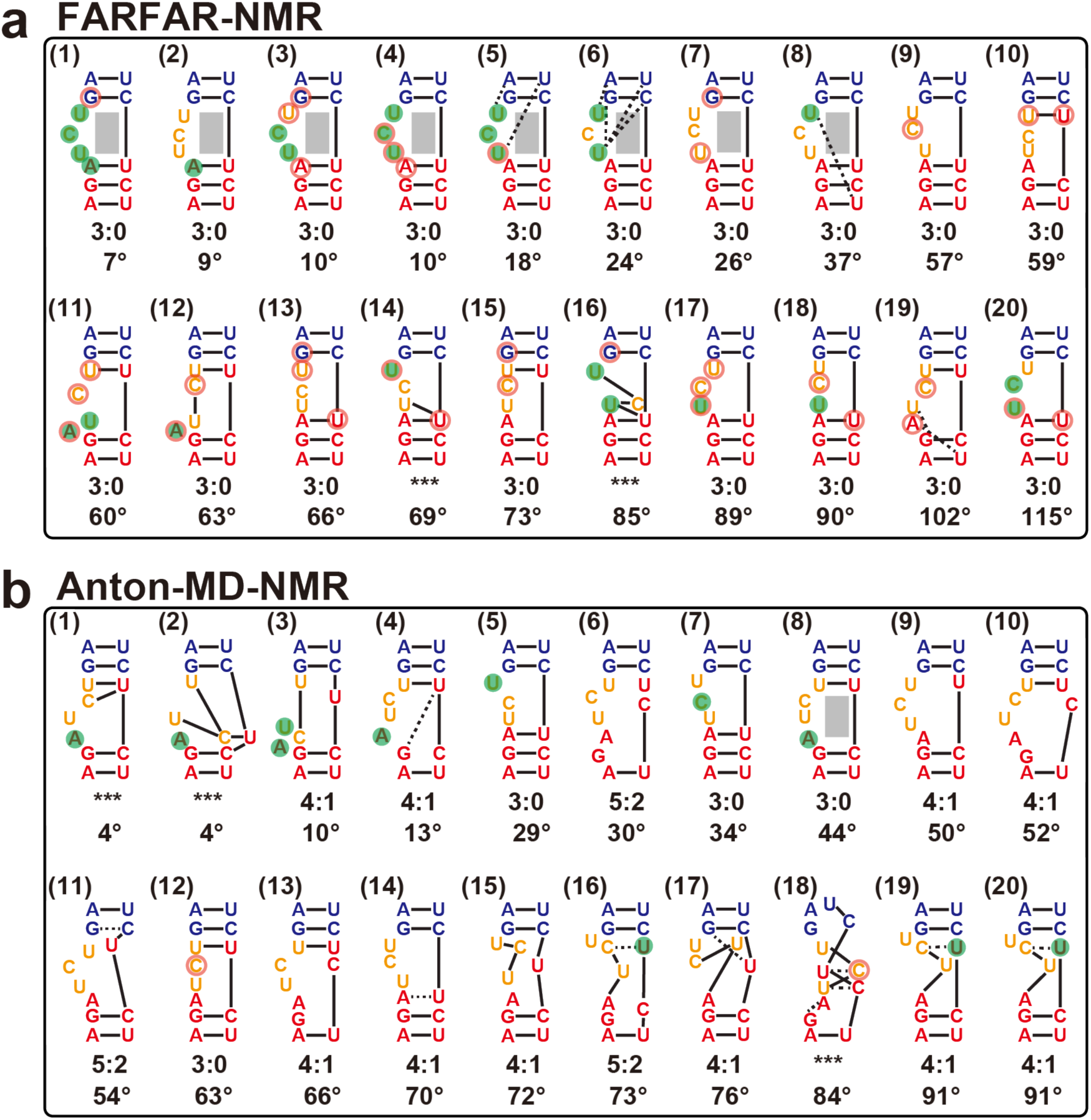
Junction topology scheme of the TAR ensembles. Junction topology in the (a) FARFAR-NMR (*N* = 20) and (b) Anton-MD-NMR (*N* = 20) ensembles. Conformers in each ensemble are sorted in increasing order of the bend angle magnitude |β_h_|, and the junction topology (Methods) as well as the |β_h_| are labeled below each conformer. Junctional residues (bulge/A22-U40/G26-C39) with C2′-*endo* sugar pucker and non-gauche+ γ (falling outside 20-100°) torsion angle are highlighted with green filled circle and orange open circle, respectively.

**Extended Data Fig. 8.**
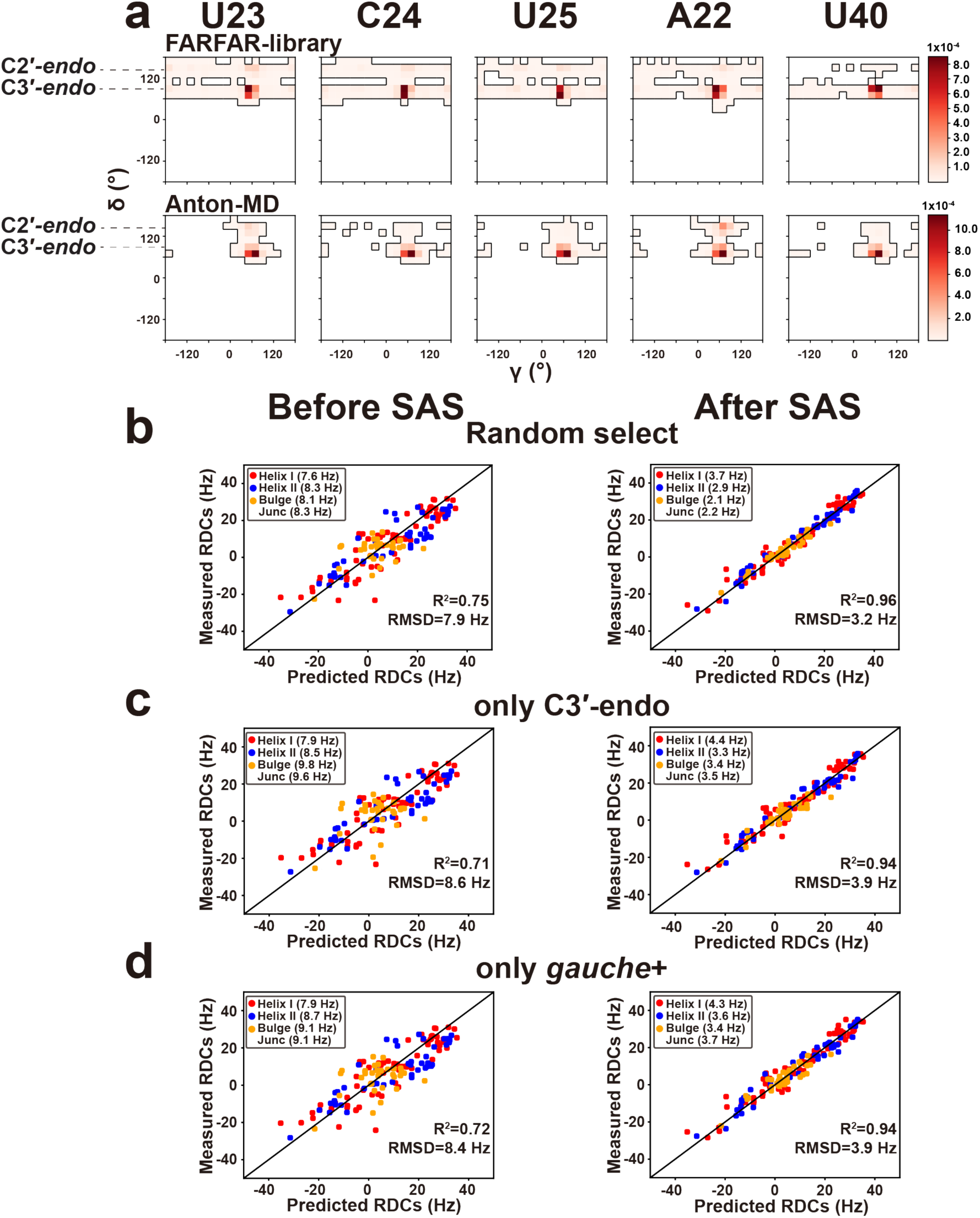
Testing the sensitivity of FARFAR-library and FARFAR-NMR ensembles to variations in local torsion angles. (a) Distribution of backbone torsion angles of the FARFAR-library and Anton-MD. 2D density map of δ versus γ of the bulge residues as well as A22 and U40, for the FARFAR-library and Anton-MD (*N* = 10,000). The bin width is 20° for all density maps. (b-d) Comparison of measured and predicted RDCs for a subset of the FARFAR-library (left, *N* = 3,000) and ensembles (right, *N* = 20) following SAS on these subset libraries. (b) Randomly selected subset from the FARFAR-library (c) a subset of the FARFAR-library with the sugar puckers of U23, C24, U25, A22 and U40 chosen to be C3′-*endo* (d) a subset of the FARFAR-library with the γ torsion angles of U23, C24, U25, A22 and U40 chosen to be gauche+ (20°<γ<100°). “Junc” denotes bulge residues as well as A22 and U40.

**Extended Data Fig. 9.**
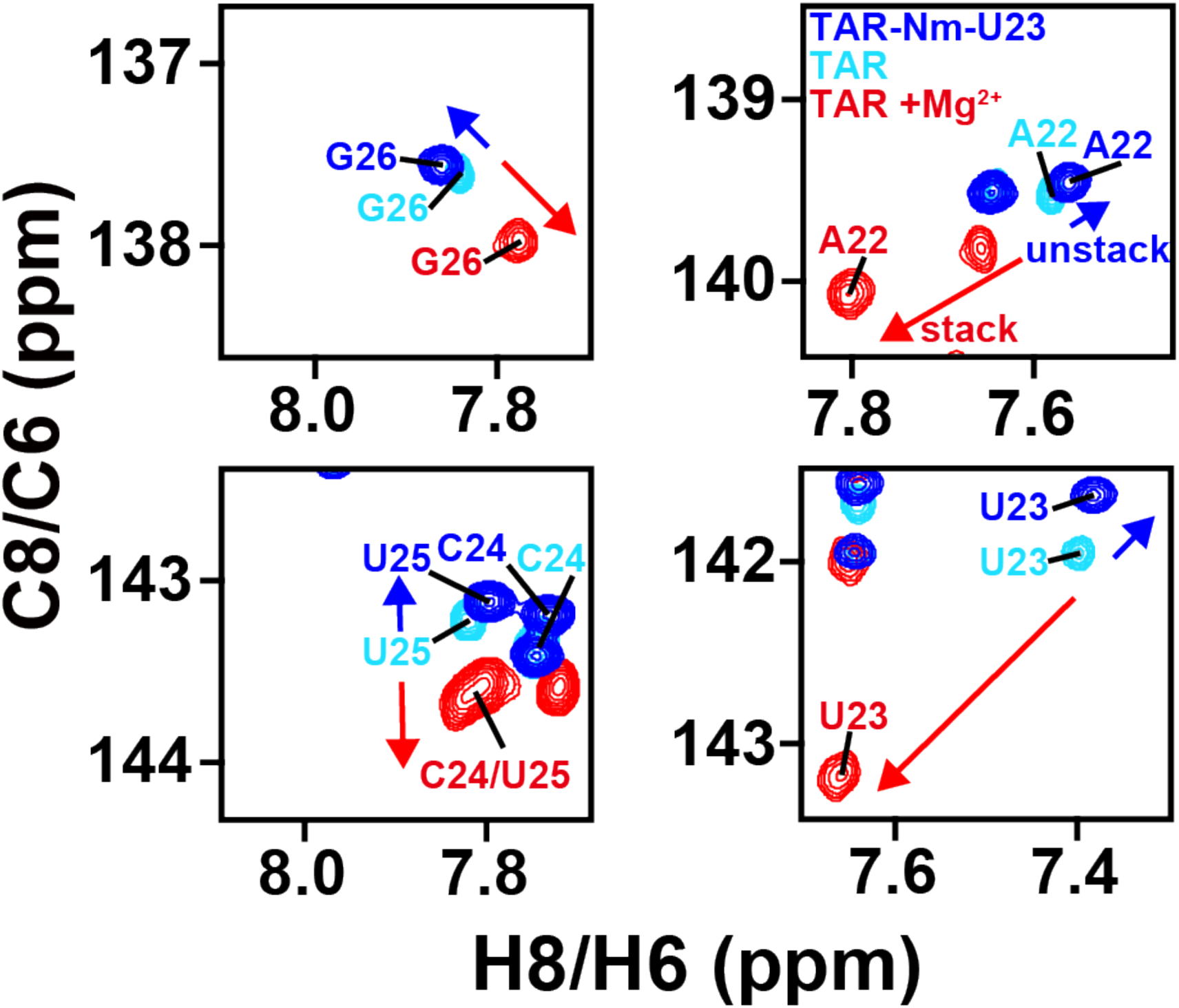
An additional Nm modified TAR showing Nm inducing unstack of TAR. Overlay of 2D [^13^C, ^1^H] HSQC NMR spectra of the aromatic region of TAR-Nm-U23 without Mg^2+^ (blue, inducing unstacking). TAR without Mg^2+^ (cyan), and TAR +Mg^2+^ (red, inducing stacking).

**Extended Data Fig. 10.**
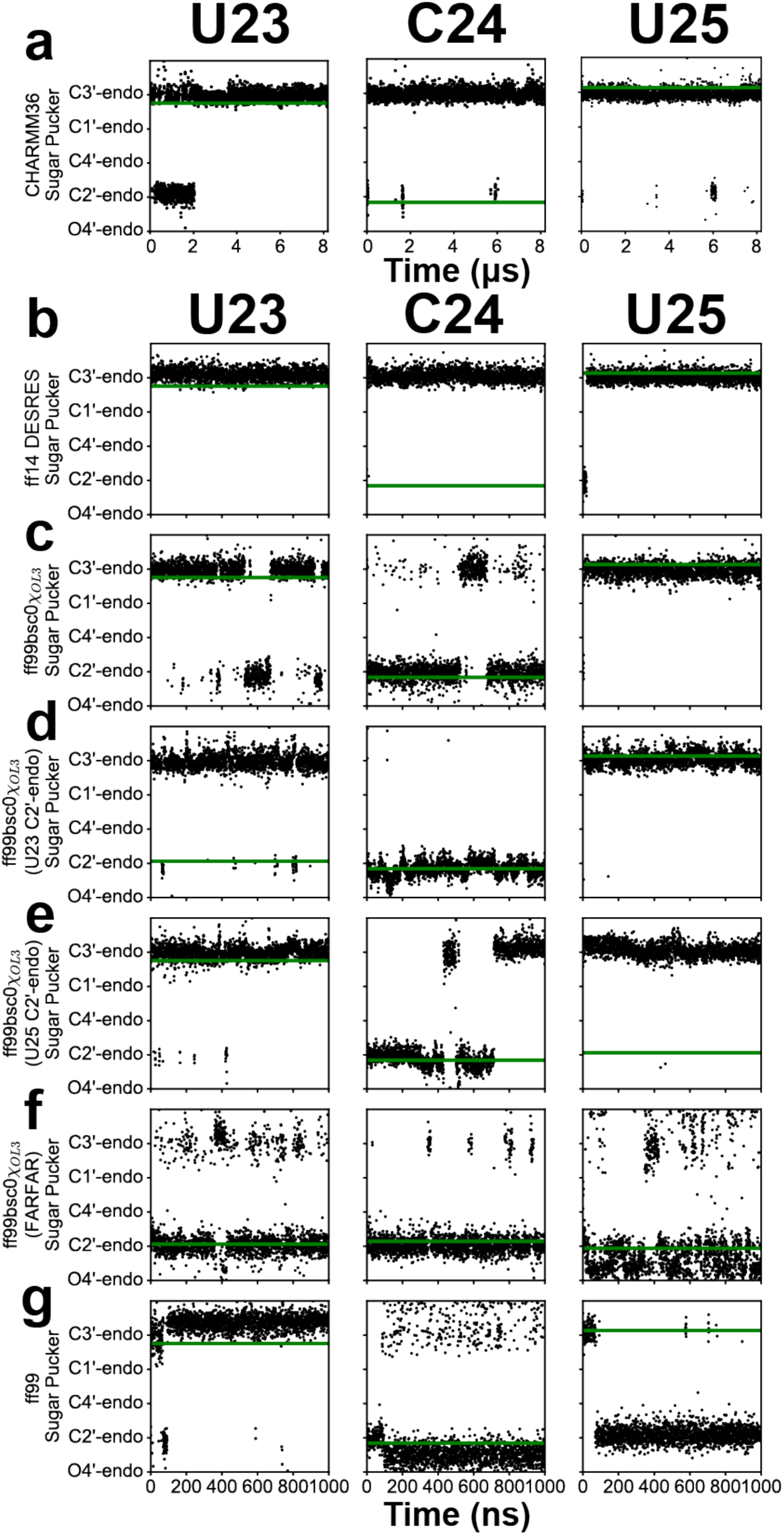
Sugar puckers of bulge nucleotides during the course of MD simulations with different force fields and starting structures. The variation of the sugar puckers of U23, C24 and U25 of TAR during the course of (a) an 8.2 µs trajectory using the Anton CHARMM36 force field and (b-g) a series of 1.0 µs trajectories using the (b) ff14 DESRES force field starting with the NOE structure (PDB: 1ANR), (c-f) ff99bsc0χOL3 force field starting with (c) the NOE structure (PDB 1ANR), (d) 1ANR with the pucker of U23 changed to C2′-*endo* (TAR^U23C2′-*endo*^), (e) 1ANR with the pucker of U25 sugar changed to C2′-*endo* (TAR^U25C2′-*endo*^), (f) a FARFAR-NMR conformer in which all bulge nucleotides are C2′-*endo* (TAR^FARFAR^), and (g) the ff99 force field. The sugar pucker of the starting structures are indicated using a green line. Persistence of a general bias towards C3′-*endo* or a tendency to maintain the sugar pucker in the initial starting conformation can be seen.

