## Supplementary Information for "An RNA dynamic ensemble at atomic resolution"

### Supplementary Discussion

FARFAR-NMR reproduces inter-helical orientations of TAR and its variants

As an initial test of the generality and limits of our new approach, we used FARFAR-NMR to generate ensembles for TAR in the presence of $Mg^{2+}$, and three TAR mutants (Extended Data Fig. 2-3), containing one (U1-TAR), two (U2-TAR) and seven (U7-TAR) bulge nucleotides in the absence and presence of $Mg^{2+}$, whose conformational dynamics have recently been characterized using NMR [1]. With the exception of U7-TAR in the absence of $Mg^{2+}$, the agreement observed between the RDCs measured for these TAR mutants (Supplementary Table 1) and values computed for the FARFAR-NMR ensembles was comparable to that obtained for TAR (Fig. 1f). The lower agreement for U7-TAR in the absence of $Mg^{2+}$ may reflect limited structural information for kinked RNAs with long bulges [1] or a need to sample a much larger number of conformations in the FARFAR calculations.

The FARFAR-NMR ensembles reproduce trends in ensemble properties observed previously for the bulge variants based on an independent analysis of RDCs and chemical shifts [1]. The average and standard deviation of the inter-helical bend angle in the FARFAR-NMR ensembles are in good agreement with values reported based on an order tensor analysis of the RDCs [1] which does not involve explicit ensemble modeling (Extended Data Fig. 3c). The distributions of the bend angle have the expected bimodal character [1] with a narrow stacked state and broader set of kinked conformations, and with $Mg^{2+}$ increasing the population of the stacked state (Extended Data Fig. 3d, e). Taken together, these results support the general applicability of FARFAR-NMR. Further refinement of these ensembles will require additional measurements of RDCs followed by evaluation using chemical shifts.

Implication of motional averaging on A-form helix

As expected, very good agreement was observed for both FARFAR and Anton-MD derived ensembles for the central Watson-Crick bps in the two helices (C19-G43, A20-U42, G21-C41, A27-U38, G28-C37), which have more rigid structures and thus should be easier to model. The slightly better agreement observed for C5′, C6 and C1′ chemical shifts for the Anton-MD-NMR as compared to the FARFAR-NMR (Extended Data Fig. 5) is presumably due to deviations from the assumed idealized static A-form geometry and/or motional averaging of the chemical shifts. Single conformers in the Anton-MD-NMR ensemble show slightly weaker agreement (overall RMSD difference < 0.22 ppm) compared to averaging over all conformers in the ensemble, including for C5′, C6 and C1′, suggesting that the better agreement in the case of Anton-MD-NMR is more likely due to neglect of motional averaging for the helices in the FARFAR-NMR ensemble.

Junctional topology dynamics of FARFAR-NMR and Anton-MD-NMR

In the FARFAR-NMR ensemble, the major topology (~75%) of the two-way junction is the canonical trinucleotide U23C24U25 bulge (3:0) (Extended Data Fig. 7a e.g. conformer (5)). However, there is also a minor topology (~25%) in which the trinucleotide bulge migrates one nucleotide down the lower stem to form a AUC bulge (Extended Data Fig. 7a e.g. conformer (10)). In contrast, in the Anton-MD-NMR ensemble for TAR, the topology of the junction varies widely (Extended Data Fig. 7b). The dominant topology (~30%) is the 4:1 internal loop lacking A22-U40 pairing and the U23C24U25 bulge, while the 3:0 internal loop with UCU bulge and a A22-U40 bp is only a minor population (~10%). The absence of base-pairing at A22-U40 is in agreement with the NMR data showing no detectable hydrogen bonds between A22 and U40. The Anton-MD-NMR topologies also include 3:0 or 4:1 internal loops (~30%) in which U25 and U40 form a U25-U40 mismatch with A22-U40 unpaired or both A22-U40 and G21-C41 unpaired, respectively. In addition, topologies with a 5:2 internal loop (~15%) in which A22-U40 as well as either G21-C41 or G26-C39 are unpaired, and those in which the entire upper helix is melted (~5 %) are also observed. Unpairing in the upper helix and at G21-C41 and G26-C39 is inconsistent with the sharp imino resonances observed for G21 and G26 [2]. These differences in pairing may help explain the better predictions of imino $^{15}N/^{1}H$ chemical shifts for the FARFAR-NMR relative to the Anton-MD-NMR ensemble (Fig. 2 and Extended Data Fig. 5). Thus, artifactual distortions in junction topology and helical base pairing in the Anton-MD ensembles appear to compensate for lack of sugar-backbone sampling to achieve the inter-helical orientations needed to satisfy the helical RDC data (Extended Data Fig. 4e-g).

### Supplementary Tables

**Supplementary Table 1. RDCs datasets used in this study**

| TAR type | Elongation | Apical loop | +/– $Mg^{2+}$ | Number of RDCs | Reference |
|---|---|---|---|---|---|
| U1-TAR | E0 | wild-type | – | 35 | 1 |
| | E0 | wild-type | + | 38 | 1 |
| U2-TAR | E0 | wild-type | – | 27 | 1 |
| | EI22 | UUCG | – | 35 | 3 |
| TAR | E0 | UUCG | – | 35 | 4 |
| | EI22 | UUCG | – | 39 | 3 |
| | EII22 | UUCG | – | 34 | 3 |
| | EI3 | UUCG | – | 35 | 5 |
| | E0 | wild-type | + | 46 | 1 |
| U7-TAR | E0 | wild-type | – | 34 | 1 |
| | E0 | wild-type | + | 36 | 1 |

**Supplementary Table 2. FARFAR input files and commands (TAR)**

<table>
<tr><td>2° structure</td><td>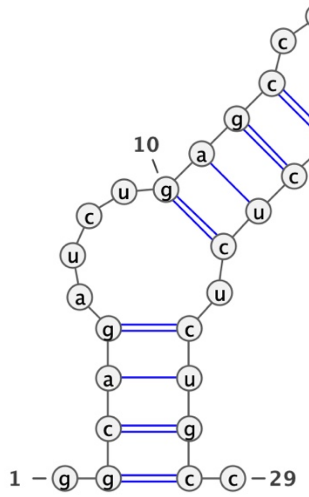<br></td></tr>
<tr><td>Input files</td><td>cat test.fasta<br>> tar<br>ggcagaucugagccugggagcucucugcc<br><br>cat test.secstruct<br>.((((....((((......)))).)))).<br>ggcagaucugagccugggagcucucugcc</td></tr>
<tr><td>Generate RNA helices</td><td>rna_helix.py -seq gcag cugc -resnum 2-5 25-28 -o helix_1.pdb -rosetta_folder ~/rosetta/main/source/cmake/build_release<br>rna_helix.py -seq gagc gcuc -resnum 10-13 20-23 -o helix_2.pdb -rosetta_folder ~/rosetta/main/source/cmake/build_release</td></tr>
<tr><td>FARFAR<br><br>run</td><td>rna_denovo -nstruct 100 -s helix_*.pdb -secstruct_file test.secstruct -fasta test.fasta -minimize_rna true</td></tr>
</table>

**Supplementary Table 3. FARFAR input files and commands (U1-TAR)**

| | |
|---|---|
| 2° structure | 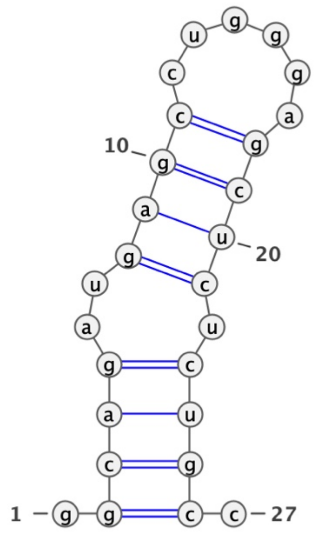 |
| Input files | *cat test.fasta*<br>*> tar*<br>*ggcagaugagccugggagcucucugcc*<br><br>*cat test.secstruct*<br>*.((((..((((......)))).)))).*<br>*ggcagaugagccugggagcucucugcc* |
| Generate<br><br>RNA helices | *rna_helix.py -seq gcag cugc -resnum 2-5 23-26 -o helix_1.pdb -rosetta_folder ~/rosetta/main/source/cmake/build_release*<br>*rna_helix.py -seq gagc gcuc -resnum 8-11 18-21 -o helix_2.pdb -rosetta_folder ~/rosetta/main/source/cmake/build_release* |
| FARFAR<br><br>run | *rna_denovo -nstruct 100 -s helix_*.pdb -secstruct_file test.secstruct -fasta test.fasta -minimize_rna true* |

**Supplementary Table 4. FARFAR input files and commands (U2-TAR)**

| | |
|---|---|
| 2° structure | 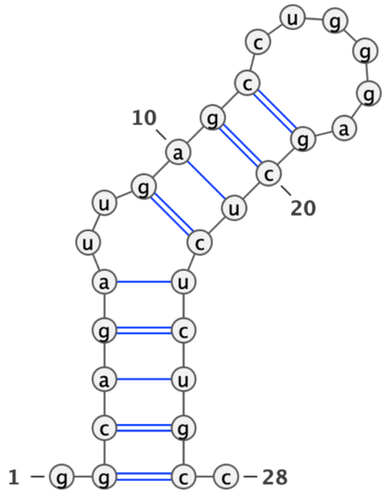  |
| Input files | cat test.fasta<br>> tar<br>ggcagauugagccugggagcucucugcc<br><br>cat test.secstruct<br>.(((((..((((......))))))))).<br>ggcagauugagccugggagcucucugcc |
| Generate<br><br>RNA helices | rna_helix.py -seq gcaga ucugc -resnum 2-6 23-27 -o helix_1.pdb -rosetta_folder ~/Rosetta/main/source/cmake/build_release<br>rna_helix.py -seq gagc gcuc -resnum 9-12 19-22 -o helix_2.pdb -rosetta_folder ~/rosetta/main/source/cmake/build_release |
| FARFAR<br><br>run | rna_denovo -nstruct 100 -s helix_*.pdb -secstruct_file test.secstruct -fasta test.fasta -minimize_rna true |

**Supplementary Table 5. FARFAR input files and commands (U7-TAR)**

| | |
|---|---|
| 2° structure | 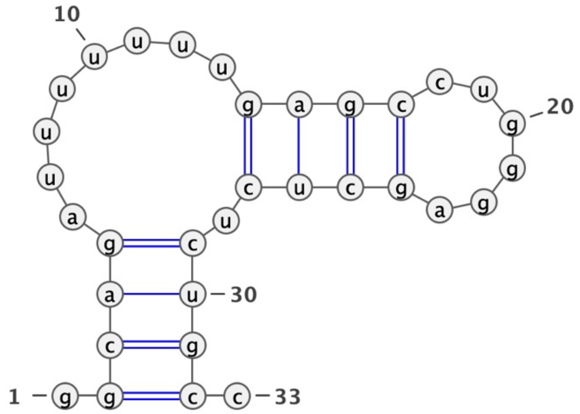 |
| Input files | `cat test.fasta`<br>`> tar`<br>`ggcagauuuuuuugagccugggagcucucugcc`<br><br>`cat test.secstruct`<br>`.((((........((((......)))).)))).`<br>`ggcagauuuuuuugagccugggagcucucugcc` |
| Generate<br><br>RNA helices | `rna_helix.py -seq gcag cugc -resnum 2-5 29-32 -o helix_1.pdb -rosetta_folder ~/rosetta/main/source/cmake/build_release`<br>`rna_helix.py -seq gagc gcuc -resnum 14-17 24-27 -o helix_2.pdb -rosetta_folder ~/rosetta/main/source/cmake/build_release` |
| FARFAR<br><br>run | `rna_denovo -nstruct 100 -s helix_*.pdb -secstruct_file test.secstruct -fasta test.fasta -minimize_rna true` |

**Supplementary Video 1.** The FARFAR-NMR atomic-resolution dynamic ensemble of HIV-1 TAR. Flipped out bulge residues are in red, flipped in bulge nucleotides are in blue, and coaxial conformations are in green.
